# dFLASH; dual FLuorescent transcription factor Activity Sensor for Histone integrated live-cell reporting and high-content screening

**DOI:** 10.1101/2023.11.21.568191

**Authors:** Timothy P. Allen, Alison E. Roennfeldt, Moganalaxmi Reckdharajkumar, Miaomiao Liu, Ronald J. Quinn, Darryl L. Russell, Daniel J. Peet, Murray L. Whitelaw, David C. Bersten

**Affiliations:** School of Biological Sciences, University of Adelaide, Adelaide, South Australia, Australia; Robinson Research Institute, University of Adelaide, South Australia, Australia; Griffith Institute for Drug Discovery, Griffith University, Brisbane, Australia; ASEAN Microbiome Nutrition Centre, National Neuroscience Institute, Singapore 169857, Singapore

## Abstract

Live-cell reporting of regulated transcription factor (TF) activity has a wide variety of applications in synthetic biology, drug discovery, and functional genomics. As a result, there is high value in the generation of versatile, sensitive, robust systems that can function across a range of cell types and be adapted toward diverse TF classes. Here we present the dual FLuorescent transcription factor Activity Sensor for Histone integrated live-cell reporting (dFLASH), a modular sensor for TF activity that can be readily integrated into cellular genomes. We demonstrate readily modified dFLASH platforms that homogenously, robustly, and specifically sense regulation of endogenous Hypoxia Inducible Factor (HIF) and Progesterone receptor (PGR) activities, as well as regulated coactivator recruitment to a synthetic DNA-Binding Domain-Activator Domain fusion proteins. The dual-colour nuclear fluorescence produced normalised dynamic live-cell TF activity sensing with facile generation of high-content screening lines, strong signal:noise ratios and reproducible screening capabilities (Z’ = 0.68-0.74). Finally, we demonstrate the utility of this platform for functional genomics applications by using CRISPRoff to modulate the HIF regulatory pathway, and for drug screening by using high content imaging in a bimodal design to isolate activators and inhibitors of the HIF pathway from a ∼1600 natural product library.

## Introduction

Cells integrate biochemical signals in a variety of ways to mediate effector function and alter gene expression. Transcription factors (TF) sit at the heart of cell signalling and gene regulatory networks, linking environment to genetic output^1,2^. TF importance is well illustrated by the consequences of their dysregulation within disease, particularly cancer where TFs drive pathogenic genetic programs^3–5^. As a result, there is widespread utility in methods to manipulate and track TF activity in basic biology and medical research, predominantly using TF responsive reporters. Recent examples include enhancer activity screening^6^ by massively parallel reporter assays, discovery and characterisation of transcription effector domains^7,8^ and CRISPR-based functional genomic screens that use reporter gene readouts to understand transcriptional regulatory networks^2,9^. Beyond the use in discovery biology TF reporters are increasingly utilised as sensors and actuators in engineered synthetic biology applications such as diagnostics and cellular therapeutics. For example, synthetic circuits that utilise either endogenous or synthetic TF responses have been exploited to engineer cellular biotherapeutics^10^. In particular, the synthetic Notch receptor (SynNotch) in which programable extracellular binding elicits synthetic TF signalling to enhance tumour-specific activation of CAR-T cells, overcome cancer immune suppression, or provide precise tumour target specificity ^11–14^.

Fluorescent reporter systems are now commonplace in many studies linking cell signalling to TF function and are particularly useful to study single cell features of gene expression, such as stochastics and heterogeneity^15^, or situations where temporal recordings are required. In addition, pooled CRISPR/Cas9 functional genomic screens rely on the ability to select distinct cell pools from a homogenous reporting parent population. Screens to select functional gene regulatory elements or interrogate chromatin context in gene activation also require robust reporting in polyclonal pools^16^. Many of the current genetically encoded reporter approaches, by nature of their design, are constrained to particular reporting methods or applications ^9,17^. For example, high content arrayed platforms are often incompatible with flow cytometry readouts and vice versa. As such there is a need to generate modular, broadly applicable platforms for robust homogenous reporting of transcription factor and molecular signalling pathways.^2^.

Here we address this by generating a versatile, high-performance sensor of signal regulated TFs. We developed a reporter platform, termed the dual FLuorescent TF Activity Sensor for Histone integrated live-cell reporting (dFLASH), that enables lentiviral mediated genomic integration of a TF responsive reporter coupled with an internal control. The well-defined hypoxic and steroid receptor signalling pathways were targeted to demonstrate that the composition of the modular dFLASH cassette is critical to robust enhancer-driven reporting. dFLASH acts as a dynamic sensor of targeted endogenous pathways as well as synthetic TF chimeras in polyclonal pools by temporal high-content imaging and flow cytometry. Routine isolation of homogenously responding reporter lines enabled robust high content image-based screening (Z’ = 0.68-0.74) for signal regulation of endogenous and synthetic TFs, as well as demonstrating utility for functional genomic investigations with CRISPRoff. Array-based temporal high content imaging with a hypoxia response element dFLASH successfully identified novel regulators of the hypoxic response pathway, illustrating the suitability of dFLASH for arrayed drug screening applications. This shows the dFLASH platform allows for intricate interrogation of signalling pathways and illustrates its value for functional gene discovery, evaluation of regulatory elements or investigations into chemical manipulation of TF regulation.

## Results

### Design of versatile dFLASH, a dual fluorescent, live cell sensor of TF activity

To fulfil the need for a modifiable fluorescent sensor cassette that can be integrated into chromatin and enable robust live-cell sensing that is adaptable for any nominated TF, applicable to high content imaging (HCI) and selection of single responding cells from polyclonal pools via image segmentation or flow cytometry (**Figure 1c**) a lentiviral construct with enhancer regulated expression of *Tomato*, followed by independent, constitutive expression of *d2EGFP* as both selectable marker and an internal control was constructed (**Figure 1a, b**). Three nuclear localisation signals (3xNLS) integrated in each fluorescent protein ensured nuclear enrichment to enable single cell identification by nuclear segmentation, with accompanying image-based quantification of normalised reporter outputs using high content image analysis, or single-cell isolation using FACS in a signal dependent or independent manner. The enhancer insertion cassette upstream of the minimal promoter driving Tomato expression is flanked by restriction sites, enabling alternative enhancer cloning (**Figure 1a**). The sensor response to endogenous signal-regulated TF pathways was first assessed by inserting a Hypoxia Inducible Factor (HIF) enhancer. HIF-1 is the master regulator of cellular adaption to low oxygen tension and has various roles in several diseases^18–20^. To mediate its transcriptional program, the HIF-1α subunit heterodimerises with Aryl Hydrocarbon Nuclear Translocator (ARNT), forming an active HIF-1 complex. At normoxia^4^, HIF-1a is post-translationally downregulated through the action of prolyl hydroxylase (PHD) enzymes and the Von Hippel Lindau (VHL) ubiquitin ligase complex^21^. Additionally, the C-terminal transactivation domain of HIF-1α undergoes asparaginyl hydroxylation mediated by Factor Inhibiting HIF (FIH), which blocks binding of transcription coactivators CBP/p300^22^. These hydroxylation processes are repressed during low oxygen conditions, enabling rapid accumulation of active HIF-1α. HIF-1α stabilisation at normoxia^4^ was artificially triggered by treating cells with the hypoxia mimetic dimethyloxalylglycine (DMOG), which inhibits PHDs and FIH, thereby inducing HIF-1α stabilisation, activity and hypoxic gene expression^23^. The well characterised regulation and disease relevance of HIF-1α made it an ideal TF target for prototype sensor development.

**Figure 1.**
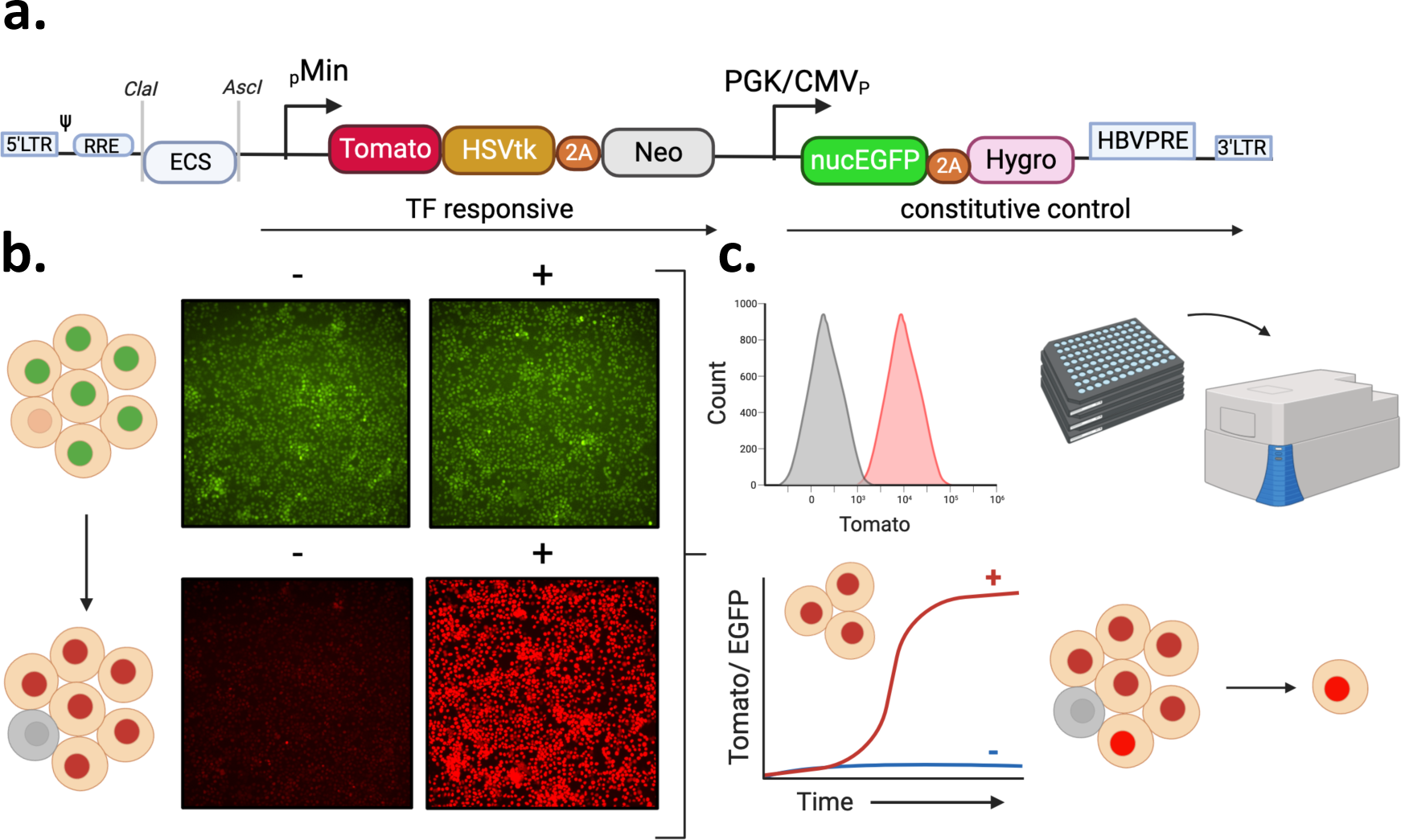
Summary of dFLASH LV-REPORT construction, utility, and validation. (**a**) The dFLASH system utilises the lentiviral LV-REPORT construct, consisting of a cis-element multiple cloning site for enhancer insertion, followed by a minimal promoter that drives a transcription factor (TF) dependent cassette that encodes three separate expression markers; a nuclear Tomato fluorophore with a 3x C-terminal nuclear localisation signal (NLS), Herpes Simplex Virus Thymidine Kinase (HSVtK) for negative selection and Neomycin resistance (Neo) for positive selection separated by a 2A self-cleaving peptide (2A). This is followed by a downstream promoter that drives an independent cassette encoding EGFP with a 3x N-terminal NLS, and a Hygromycin resistance selection marker separated by a 2A peptide. (**b**) This design allows for initial identification of the EGFP fluorophore in nuclei, independent of signal. Expression of the Tomato fluorophore is highly upregulated in a signal-dependent manner. Images shown are monoclonal HEK293T dFLASH-HIF cells. Populations were treated for 48 hours ±DMOG induction of HIF-1α and imaged by HCI. (**c**) This system can be adapted to a range of different applications. This includes (clockwise) flow cytometry, arrayed screening in a high throughput setting with high content imaging, isolation of highly responsive clones or single cells from a heterogenous population or temporal imaging of pooled or individual cells over time.

### Optimisation of dFLASH sensors

Initially, we tested FLASH constructs with repeats of hypoxia response element (HRE) containing enhancers (RCGTG)^24^ from endogenous target genes (HRE-FLASH), controlling expression of either nuclear mono (m) or tandem dimer (td)Tomato and observed no DMOG induced Tomato expression in stable HEK293T cell lines (mnucTomato or tdnucTomato, **Supp Figure 1a,b**). Given the HIF response element has been validated previously^24^, the response to HIF-1α was optimised by altering the reporter design, all of which utilised the smaller mnucTomato (vs tdnucTomato) to contain transgene size. We hypothesised that transgene silencing, chromosomal site-specific effects or promoter enhancer coupling/interference may result in poor signal induced reporter activity observed in initial construct designs. As such we optimised the downstream promoter, the reporter composition and incorporated a 3xNLS d2EGFP internal control from the constitutive promoter to monitor chromosomal effects and transgene silencing.

Dual FLASH (dFLASH) variants incorporated three variations of the downstream promoter (EF1a, PGK and PGK/CMV) driving *3xNLS EGFP (nucEGFP)* and 2A peptide linked hygromycin (detailed in **Supp Figure 1c**) in combination with alternate reporter transgenes that it expressed mnucTomato alone, or mnucTomato-Herpes Simplex Virus Thymidine Kinase (HSVtK)-2A-Neomycin resistance (Neo). Stable HEK293T and HepG2 HRE-dFLASH cells lines with these backbones were generated by lentiviral transduction and hygromycin selection. The reporter efficacy of dFLASH variant cell lines was subsequently monitored by high content imaging 48 hours after DMOG induction (**Supp Figure 1d, e)**. The downstream composite PGK/CMV or PGK promoters, enabled the strong DMOG induced Tomato or Tomato/GFP expression dramatically outperforming EF1a (**Figure 1b** and **Supp Figure 1d**). The composite PGK/CMV provided bright, constitutive nucEGFP expression in both HepG2 and HEK293T cells which was unchanged by DMOG, whereas nucEGFP controlled by the PGK promoter was modestly increased (∼2.5 fold) by DMOG (**Supp Figure 1e**). Substitution of the mnucTomato with the longer mnucTomato-HSVtK-Neo reporter had no effect on DMOG induced reporter induction in EF1a containing HRE-dFLASH cells, still failing to induce tomato expression (**Supp Figure 1f**). CMV/PGK containing dFLASH sensors maintained DMOG induction when either the mnucTomato or the mnucTomato/HSVtK/Neo reporters were utilised (**Supp Figure 1g, h**) although mnucTomato without HSVtK and Neo produced lower absolute mnucTomato fluorescence and a smaller percentage of cells responding to DMOG, albeit with lower background. Taken together these findings indicate that certain backbone compositions prevented or enabled robust activation of the enhancer driven cassette, similar to the suppression of an upstream promoter by a downstream, contiguous promoter previously described^25,26^ suggesting that the 3’ EF1a promoter results in poorly functioning multi-cistronic synthetic reporter designs^27^. Consequently, the PGK/CMV backbone and the mnucTomato/HSVtK/Neo reporter from **Supp Figure 1** was chosen as the optimised reporter design (HRE-dFLASH). To confirm that the HRE element was conferring HIF specificity, a no response element dFLASH construct in HEK293T cells treated with DMOG produced no change in either mnucTomato or nucEGFP compared to vehicle-treated populations (**Supp Figure 2a**). This result, together with the robust induction in response to DMOG (**Figure 2D**, **Supp Figure 1f**, **1h**), confirms HIF enhancer driven reporter to respond robustly to induction of the HIF pathway (subsequently labelled dFLASH-HIF).

**Figure 2.**
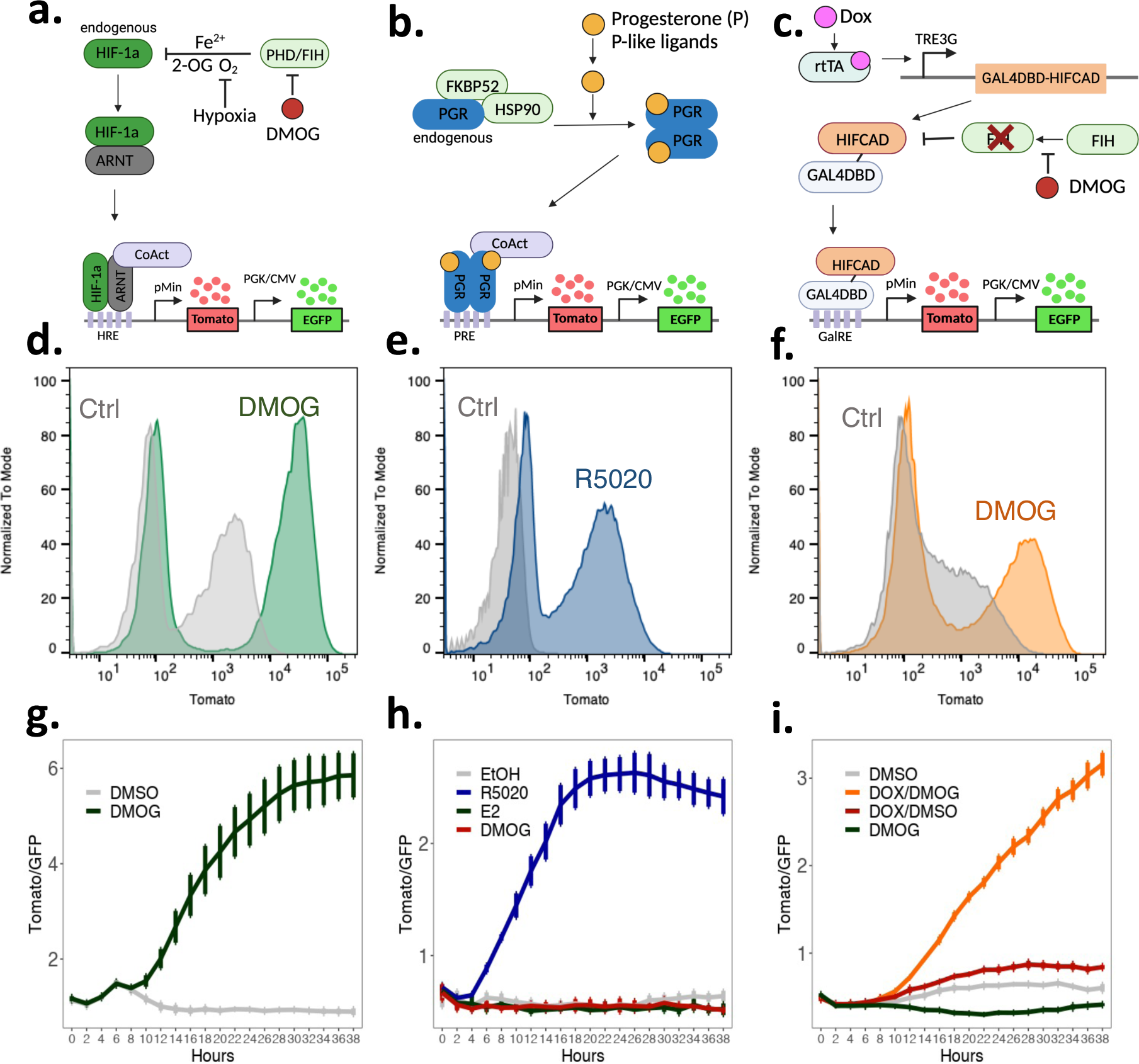
dFLASH provides sensitive readouts to three distinct TF pathways. (**a-c**) Three distinct enhancer elements enabling targeting of three different signalling aspects. (**a**) Hypoxic response elements (HRE) provide a read out for HIF-1α activation; (**b**) Progesterone response elements (PRE) derived from progesterone receptor target genes facilitate reporting of progestin signaling; (**c**) Gal4 response elements (GalRE) enable targeting of synthetic transcription factors to dFLASH such as a GAL4DBD-HIFCAD fusion protein that provides a FIH-dependent reporter response. (**d-f**) Flow cytometry histograms showing Tomato expression following 48 hr treatments of the indicated dFLASH polyclonal reporter cells (**d**) HEK293T; 1mM DMOG or 0.1% DMSO (Ctrl), (**e**) T47D; 100nM R5020 or Ethanol (Ctrl), (**f**) HEK293T; 1μg/mL Doxycycline (Dox) and 1mM DMOG or Dox and 0.1% DMSO (Ctrl). (**g-i**) Reporter populations as in **d-f** were temporally imaged for 38 hours using HCI directly after treatment with (**g**) 0.5mM DMOG or 0.1% DMSO, (4 replicates) (**h**) 100nM R5020, 35nM E2, 0.5mM DMOG or 0.1% Ethanol (EtOH) (4 replicates), (**i**) 0.1% DMSO, 1mM DMOG, 100ng/mL Dox and 0.1% DMSO, or 100ng/mL Dox and 1mM DMOG (4 replicates).

To validate the high inducibility and nucEGFP independence of dFLASH was not specific to the HIF pathway, we generated a Gal4 responsive dFLASH construct (Gal4RE-dFLASH), using Gal4 responsive enhancers^22,28^. HEK293T cells were transduced with Gal4RE-dFLASH and a dox-inducible expression system to express synthetic Gal4DBDtransactivation domain fusion protein. To evaluate Gal4RE-dFLASH we expressed Gal4DBD fused with a compact VPR (miniVPR), a strong transcriptional activator^29^ (**Supp Figure 2b, 3a-c**). We observed ∼25% of the polyclonal population was highly responsive to doxycycline treatment (**Supp Figure 2b**), with a ∼14-fold change in Tomato expression relative to nucEGFP by HCI (**Supp Figure 3c**) demonstrating our optimised dFLASH backbone underpins a versatile reporting platform.

**Figure 3.**
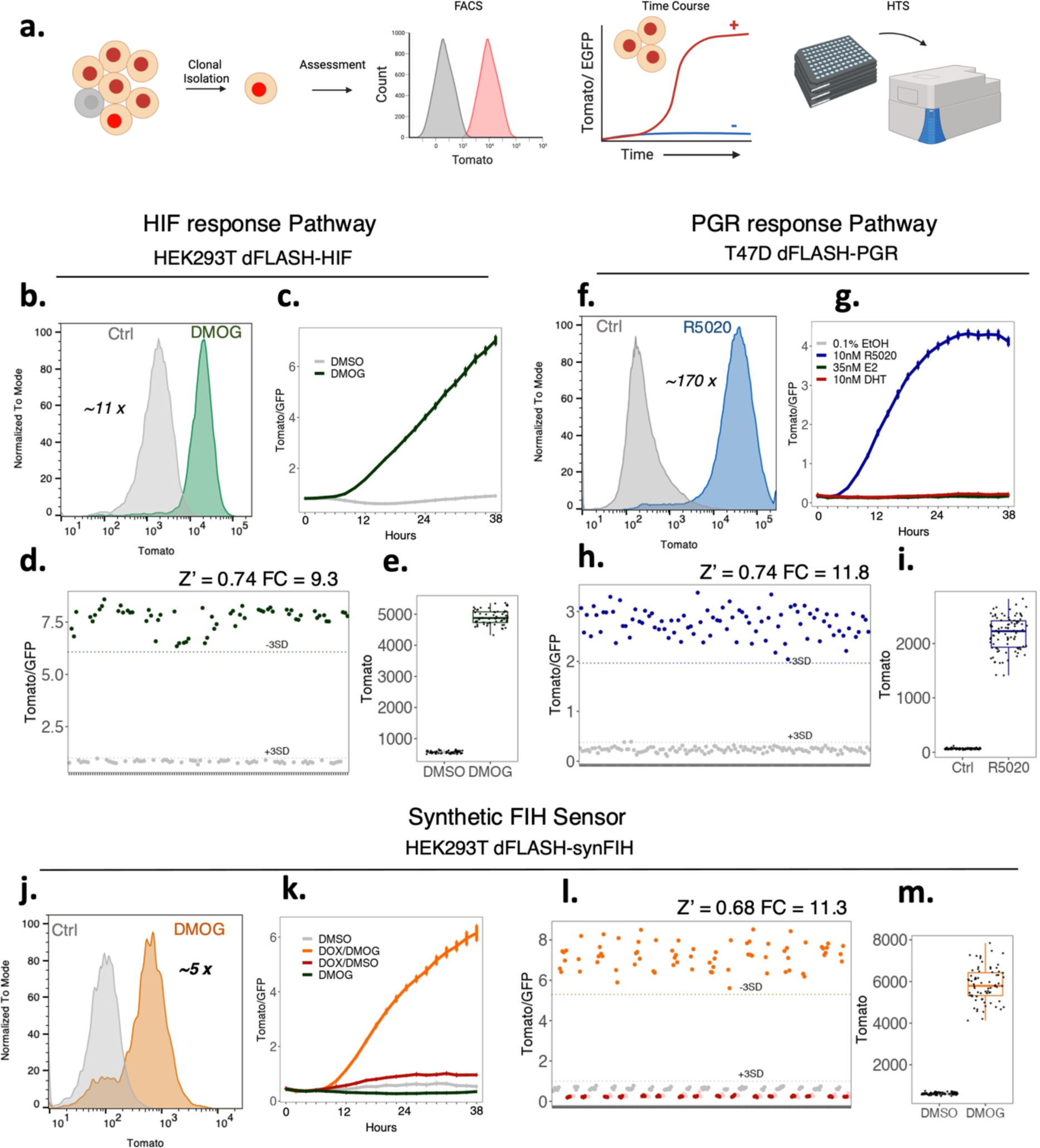
Derivation of robust, screen-ready dFLASH clonal lines. (**a**) Schematic for derivation and assessment of robustness for clonal lines of (**b-e**) HEK239T dFLASH-HIF (mcdFLASH-HIF), (**f-i**) T47D dFLASH-PGR (mcdFLASH-PGR) and (**j-m**) HEK293T dFLASH-synFIH (mcdFLASH-synFIH) were analysed by flow cytometry, temporal HCI over 38 hours and for inter-plate robustness by mock multi-plate high throughput screening with HCI. (**b-e**) mcdFLASH-HIF was (**b**) treated with DMOG for 48 hours and assessed for Tomato induction by flow cytometry relative to vehicle controls with fold change between populations displayed and (**c**) treated with vehicle or 0.5mM DMOG and imaged every 2 hours for 38 hours by HCI (mean ±sem, 8 replicates). (**d-e**) mcdFLASH-HIF was treated for 48 hours with 1mM DMOG or vehicle (6 replicates/plate, n = 10 plates) by HCI in a high throughput screening setting (HTS-HCI) for (**d**) normalised dFLASH expression and (**e**) Tomato MFI alone. (**f – i**) T47D mcdFLASH-PGR was (**f**) assessed after 48 hours of treatment with 100nM R5020 by flow cytometry for Tomato induction and (**g**) treated with 10nM R5020, 35nM E2, 10nM DHT and vehicle then imaged every 2 hours for 38 hours by temporal HCI for normalised dFLASH expression (mean ±sem, 8 replicates). (**h-i)** T47D mcdFLASH-PGR was assessed by HTS-HCI at 48 hours (24 replicates/plate, n = 5 plates) for (**h**) dFLASH normalised expression and (**i**) Tomato MFI alone. (**j**) HEK293T dFLASH-synFIH was assessed, with 200ng/mL and or Dox and 1mM DMOG by flow cytometry for dFLASH Tomato induction (**k**) mcdFLASH-synFIH was treated with 100ng/mL Dox, 1mM DMOG and relevant vehicle controls and assessed for reporter induction by temporal HCI (mean ±sem 4 replicates). (**l-m**) mcdFLASH-synFIH cells were treated with 200ng/mL Dox (grey), 1mM DMOG (red), vehicle (pink) or Dox and DMOG (orange) and assessed by HTS-HCI after 48 hours (24 replicates/plate, n = 3 plates) for (**l**) normalised dFLASH expression or (**m**) Tomato MFI induction between Dox and Dox and DMOG treated populations. Dashed lines represent 3SD from relevant vehicle (+3SD) or requisite ligand treated population (−3SD). Fold change for flow cytometry and HTS-HCI (FC) is displayed. Z’ was calculated from all analysed plates by HTS-HCI. Z’ for all plates analysed was > 0.5.

### dFLASH senses functionally distinct TF activation pathways

Following the success in utilising dFLASH to respond to synthetic transcription factor and HIF signalling, we explored the broader applicability of this system to sense other TF activation pathways. We chose the Progesterone Receptor (PGR), a member of the 3-Ketosteriod receptor family that includes the Androgen, Glucocorticoid and Mineralocorticoid receptors, as a functionally distinct TF pathway with dose-dependent responsiveness to progestin steroids to assess the adaptability of dFLASH performance. Keto-steroid receptors act through a well-described mechanism which requires direct ligand binding to initiate homodimerization via their Zinc finger DNA binding domains, followed by binding to palindromic DNA consensus sequences. PGR is the primary target of progesterone (P4, or a structural mimic R5020) and has highly context dependent roles in reproduction depending on tissue type^30,31,32^. We inserted PGR-target gene enhancer sequences containing the canonical NR3C motif (ACANNNTGT^31^) into dFLASH, conferring specificity to the ketosteroid receptor family to generate PRE-dFLASH (**Figure 2b**, see **Methods**).

A chimeric TF system was also established with Gal4DBD fusion proteins to create a synthetic reporter to sense the enzymatic activity of oxygen sensor Factor Inhibiting HIF (FIH). This sensor system termed SynFIH for its ability to synthetically sense FIH activity contained Gal4DBD-HIFCAD fusion protein expressed in a doxycycline-dependent manner, in cells harbouring stably integrated Gal4RE-dFLASH. FIH blocks HIF transactivation through hydroxylation of a conserved asparagine in the HIF-1α C-terminal transactivation domain (HIFCAD), preventing recruitment of the CBP/p300 co-activator complex^22^. As FIH is a member of the 2-oxoglutarate dioxygenase family, like the PHDs which regulate HIF post-translationally, it is inhibited by DMOG (**Figure 2C**), allowing induction of SynFIH-dFLASH upon joint Dox and DMOG signalling (**Supp Figure 3d,3e**). dFLASH-based sensors for PGR and Gal4DBD-HIFCAD generated in the optimised backbone used for dFLASH-HIF (**Figure 2a-c**). For the PGR sensor we transduced T47D cells with PRE-dFLASH, as these have high endogenous PGR expression, while for the FIH-dependent system we generated HEK293T cells with Gal4RE-dFLASH and the GAL4DBD-HIFCAD system (dFLASH-synFIH).

Stable polyclonal cell populations were treated with their requisite chemical regulators and reporter responses analysed by either flow cytometry or temporal imaging using HCI at 2hr intervals for 38 hours (**Figure 2**). Flow cytometry revealed all three systems contain a population that strongly induced nucTomato and maintained nucEGFP (**Supp Figure 2**). In HEK293T cells, ∼20% of dFLASH-synFIH and ∼50% of dFLASH-HIF population induced Tomato fluorescence substantially relative to untreated controls (**Figure 2d**, **Figure 2f**). The ∼20% reporter response to inhibition of FIH activity by DMOG (**Supp Figure 2e**, **Figure 2f**) is comparable with what was observed for GalRE-dFLASH response to Gal4DBD-miniVPR expression after equivalent selection (**Supp Figure 2b**). The PGR reporter in T47D cells showed ∼50% of the population substantively induced Tomato (**Figure 2e, Supp Figure 2d)**. The presence of considerable responsive populations for FIH, PGR, and HIF sensors, reflected in the histograms of the EGFP positive cells (**Figure 2d-f**) indicated that isolation of a highly responsive clone or subpopulations can be readily achievable for a range of transcription response types. Importantly, the induction of dFLASH-synFIH by Dox/DMOG co-treatment was ablated and displayed high basal Tomato levels in FIH knockout dFLASH-synFIH cells (**Supp Figure 3e**), indicating that the dFLASH-synFIH specifically senses FIH enzymatic activity.

All dFLASH systems showed consistent signal-dependent increases in reporter activity out to 38 hours by temporal HCI enabling polyclonal populations of dFLASH to track TF activity (**Figure 2g-i**). PRE-dFLASH was more rapidly responsive to R5020 ligand induction (∼6 hours, **Figure 2h**) than dFLASH-HIF and dFLASH-synFIH to DMOG or Dox/DMOG treatment, respectively (∼10 hours, **Figure 2g, i**). Treatment of PRE-dFLASH with estrogen (E2), which activates the closely related Estrogen Receptor facilitating binding to distinct consensus DNA sites to the PGR, or the hypoxia pathway mimetic DMOG, failed to produce a response on PRE-dFLASH (**Figure 2h**). This indicates that the PRE enhancer element is selective for the ketosteroid receptor family (also see below), and that enhancer composition facilitates pathway specificity. We also observed a signal-dependent change in EGFP expression by flow cytometry in the T47D PRE-dFLASH reporter cells (**Supp Figure 2g**) but did not observe a significant change in EGFP expression for HEK293T or HEPG2 dFLASH-HIF (**Supp Figure 1c, Supp Figure 2c**) or in HEK293T dFLASH-synFIH cells (**Supp Figure 2h**), with only a small change with Gal4RE-dFLASH with Gal4DBD-miniVPR (**Supp Figure 2b**). While this change in T47D cells was not detected in the other cellular contexts (see below), it highlights that care needs to be taken in confirming the utility of the constitutive nucEGFP as an internal control in certain scenarios.

### Monoclonal dFLASH cell lines confer robust screening potential in live cells

The observed heterogenous expression of dFLASH within polyclonal cell pools is useful in many assay contexts but reduces efficiency in arrayed high content screening experiments and incompatible with pooled isolation of loss of function regulators. Therefore, monoclonal HEK293T and HepG2 dFLASH-HIF, T47D and BT474 PRE-dFLASH and HEK293T dFLASH-synFIH cell lines were derived to increase reliability of induction, as well as consistency and homogeneity of reporting (**Figure 3, Supp Figure 4**). The isolated mcdFLASH-synFIH and mcdFLASH-HIF lines also demonstrated constitutive signal insensitive nucEGFP expression (**Supp Figure 4a,b,i**). While the T47D PRE-mcdFLASH showed a small increase in nucEGFP in response to R5020, this did not preclude the use in normalisation of high content imaging experiments (see below).

**Figure 4.**
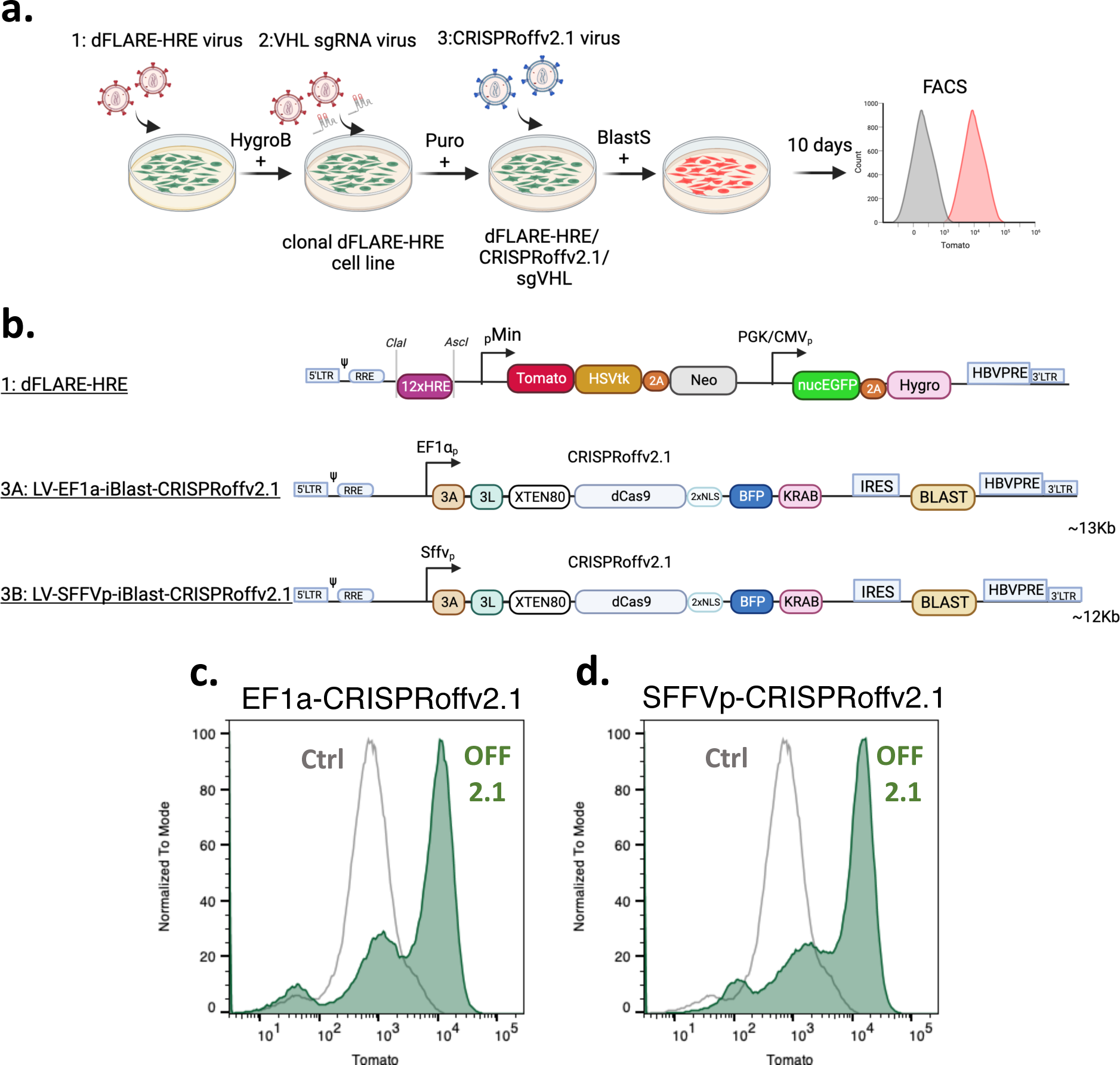
Near homogenous activation of mcdFLASH-HIF by CRISPRoff knockdown of VHL. (**a**) Clonal (1) mcdFLASH-HIF lines derived post-hygromycin (HygroB) selection were transduced first with the (2) sgRNA vector targeting *VHL* transcriptional start site, followed by puromycin selection (Puro). This pool was subsequently transduced by the (3) CRISPRoffv2.1 virus and selected with blasticidin (BlastS) prior to flow cytometry (on day 5 and 10 post Blasticidin selection). (**b**) The (1) dFLASH vector with the HRE enhancer was transduced as were 2 variants of the CRISPRoffv2.1 vector with either (**3A**) EF1α promoter or (**3B**) SFFV promoter driving the dCas9 expression cassette. (**c, d**) Flow cytometry for dFLASH-HIF induction in response to the CRISPRoffv2.1 VHL knockdown relative to parental line (Ctrl) with (**c**) EF1a or (**d**) SFFV expression constructs after 10 days of selection.

No change in EGFP in BT474 PRE-mcdFLASH cells indicates that strong transactivation leading to promoter read through or cell-type specific effects may be at play. Flow cytometry of monoclonal dFLASH cell lines with their cognate ligand inducers (DMOG (**Figure 3b**), R5020 (**Figure 3f**) or Dox/DMOG (**Figure 3j**)) revealed robust homogeneous induction of mnucTomato in all cell lines. Using temporal high content imaging we also found that clonally derived lines displayed similar signal induced kinetics as the polyclonal reporters although displayed higher signal to noise and increased consistency (**Figure 3, Supp Figure 4i**). Using physiologically relevant concentrations of steroids or steroid analogs (10nM-35nM), the PRE-mcdFLASH lines selectively respond to R5020 (10nM) not E2 (35nM), DHT (10nM), Dexamethasone (Dex, 10nM) or Retinoic acid (RA, 10nM) (**Figure 3g, Supp Figure 4i**). In addition, dose response curves of R5020 mediated Tomato induction indicate that PRE-mcdFLASH line responds to R5020 with an EC_50_ ∼200pM, in agreement with orthogonal methods^33^ (**Supp Figure 4g, h**). This suggests that the PRE-mcdFLASH responds sensitively and selectively to PGR selective agonist R5020, with the potential for high-content screening for modulators of *PGR* activity. As such, we term this line mcdFLASH-PGR from herein, for its specific ability to report on PGR activity at physiological steroid concentrations.

The temporal HCI of populations (**Figure 2 and Figure 3**) were imaged every 2hrs and do not inherently provide single-cell temporal dynamics of transcriptional responses. Using clonally derived mcdFLASH-PGR or mcdFLASH-HIF lines we also imaged transcriptional responses to R5020 or DMOG, respectively every 15mins (**Supp Video 1 and 2**). High temporal resolution imaging has the potential to monitor transcriptional dynamics in single cells, facilitated by the dual fluorescent nature of dFLASH. Taken together this indicates that clonal lines display improved signal to noise and assay consistency, possibly enabling high content screening experiments.

Typically, high-content screening experiments require high in-plate and across plate consistency, therefore we evaluated mcdFLASH lines (HIF-1α, PGR, FIH) across multiple plates and replicates. System robustness was quantified with the Z’ metric^34^ accounting for fold induction and variability between minimal and maximal dFLASH outputs. Signal induced mnucTomato fluorescence across replicates from independent plates was highly consistent (Z’ 0.68-0.74) and robust (9.3-11.8 fold, **Figure 3 d, h, l**) the signal induced changes in activity for mcdFLASH-HIF and mcdFLASH-FIH were driven by increased mnucTomato, with minimal changes in nucEGFP (**Figures 3e and 3m**). Despite the changes previously observed in nucEGFP mcdFLASH-PGR in T47D cells provided equivalent reporter to the other systems, (**Figure 3h, i**) as a result, monoclonal mcdFLASH cell lines represent excellent high-throughput screening systems routinely achieving Z’ scores > 0.5. Importantly, the induction of the mcdFLASH lines (HEK293T and HepG2 mcdFLASH-HIF, T47D mcdFLASH-PGR and HEK293T mcdFLASH-SynFIH) remained stable over extended passaging (months), enabling protracted large screening applications.

### dFLASH-HIF CRISPR-perturbations of the HIF pathway

The robust signal window and high Z’ score of mcdFLASH-HIF cell line, coupled with facile analysis by flow cytometry and HCI, indicates that the reporter system is amenable to functional genomic screening. We utilised the recently developed CRISPRoffv2.1 system^35^ to stably repress expression of VHL, which mediates post-translational downregulation of the HIF-1α pathway ^36,37^. We generated stable mcdFLASH-HIF cells expressing a guide targeting the VHL promoter and subsequently introduced CRISPRoffv2.1 from either a lentivirus driven by an EF1a or SFFV promoter (**Figure 4a, b**). Cells were then analysed by flow cytometry 5-or 10-days post selection to determine if measurable induction of mcdFLASH-HIF reporter was modulated by VHL knockdown under normoxic conditions (**Supp Figure 6**, **Figure 4c, 4d)**. As expected, mcdFLASH-HIF/sgVHL cells expressing CRISPRoffv2.1 from either promoter induced the mcdFLASH-HIF reporter in ∼35% by 5 days and the majority of cells (∼60%) by 10 days as compared to parental cells. Demonstration that mcdFLASH-HIF is responsive to CRISPRi/off perturbations of key regulators of the HIF pathway illustrates the potential for the dFLASH platform to provide a readout for CRISPR screens at-scale in a larger format including genome-wide screens.

### dFLASH facilitates bimodal screening for small molecule discovery

Manipulation of the HIF pathway is an attractive target in several disease states, such as in chronic anaemia^38^ and ischemic disease^39^ where its promotion of cell adaption and survival during limiting oxygen is desired. Conversely, within certain cancer subtypes^40,41^ HIF signalling is detrimental and promotes tumorigenesis. Therapeutic agents for activation of HIF-α signalling through targeting HIF-α regulators were initially discovered using *in vitro* assays. However, clinically effective inhibitors of HIF-1α signalling are yet to be discovered^42^. The biological roles for HIF-1α and closely related isoform HIF-2α, which share the same canonical control pathway, can be disparate or opposing in different disease contexts requiring isoform selectivity for therapeutic intervention^43^. To validate that HIF-1α is the sole isoform regulating mcdFLASH-HIF in HEK293T cells^44^ tandem HA-3xFLAG epitope tags were knocked in to the endogenous HIF-1α and HIF-2α C-termini allowing directly comparison by immunoblot^45^ and confirmed HIF1a is predominant isoform (**Supp Figure 5a**). Furthermore, there was no change in DMOG induced mnucTomato expression in HEK293T mcdFLASH-HIF cells when co-treated for up to 72 hours with the selective HIF2a inhibitor PT-2385 (**Supp Figure 5b)**, consistent with the minimal detection of HIF-2α via immunoblot. This confirmed that our HEK293T dFLASH-HIF cell line specifically reports on HIF-1 activity and not HIF-2, indicating that it may be useful for identification of drugs targeting the HIF-1α pathway.

dFLASH-HIF facilitates multiple measurements across different treatment regimens and time points, enabling capture of periodic potentiated and attenuated HIF signalling during a single experiment. Having validated the robust, consistent nature of mcdFLASH-HIF, we exploited its temporal responsiveness for small molecule discovery of activators or inhibitors of HIF-1α signalling in a single, bimodal screening protocol. To test this bimodal design, we utilised a natural product library of 1595 compounds containing structures that were unlikely to have been screened against HIF-1α prior. We first evaluated library compounds for ability to activate the reporter after treatment for 36 hours (**Figure 5a**) or 24 hours (**Figure 5d**). The selection of two different screening time points was to minimise any potential toxic effects of compounds at the later time points. Consistency of compound activity between the two screens was assessed by Pearson correlations (**Supp Figure 7i**, R = 0.79, p < 2.2×10^−16^). Lead compounds were identified by their ability to increase mnucTomato/nucEGFP (**Figure 5b, c**) and mnucTomato MFI more than 2SD compared with vehicle controls, with less than 2SD decrease in nucEGFP (21/1595 compounds (1.3%) each expt; **Supp Figure 7a, e**) and an FDR adjusted P score <0.01 across both screens (3/1595 (0.18%) compounds; **Supp Figure 7b, f**). After imaging of reporter fluorescence to determine these compound’s ability to activate HIF-1α we then treated the cells with 1mM DMOG and imaged after a further 36-hour (**Figure 5c**) and 24-hour (**Figure 5f**) period. Again, consistency of compound activity was assessed by Person correlation (**Supp Figure 7j, F**, R = 0.62, p < 2.2×10^−16^). Lead compounds were defined as those exhibiting a decrease in mnucTomato MFI >2SD from DMOG-treated controls in each screen without changing nucEGFP >2SD relative to the DMOG-treated controls (26/1595 compounds (1.3%) (36hr treatment) and 13/1595 compounds (<1%) (24hr treatment); **Supp Figure 7c, g**), and decrease in mnucTomato/nucEGFP >2SD with an FDR adjusted P score < 0.01 (3/1595 compounds (0.18%) across both expt; **Supp Figure 7d, h**).

**Figure 5.**
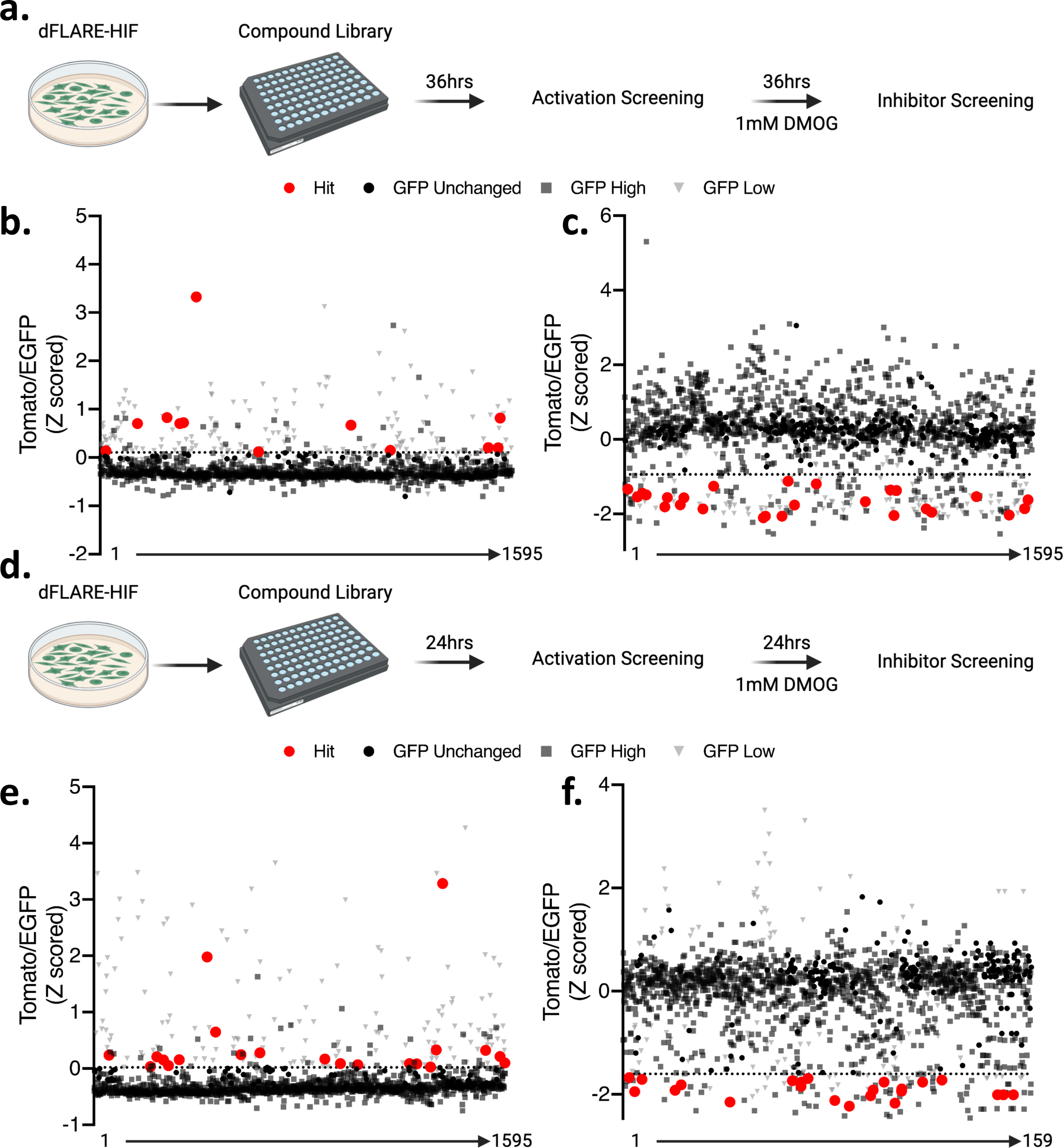
Bimodal small molecule screening of the HIF signalling pathway with dFLASH-HIF identifies positive and negative regulators. (**a**) HEK293T mcdFLASH-HIF cells were treated with a 1595 compound library and incubated for 36 hours prior to (**b**) the first round of HCI normalised dFLASH activity. Compounds that changed EGFP >±2SD are shown in grey and excluded as hits. Compounds that increase Tomato/EGFP >2SD from the vehicle controls (dashed line) are highlighted in red. After the activation screen, the compound wells were then treated with 1mM DMOG for 36 hours prior to the second round of HCI. Compounds that decreased dFLASH activity greater than 2SD from DMOG controls (dashed line) are shown in red. Compounds that changed EGFP >±2SD are shown in grey and excluded as hits. Normalised dFLASH output (Z scoring) for all analysed wells. (**d-f**) The screening protocol of (**a-c**) was repeated using 24 hr points for HCI.

### dFLASH identified novel and known compounds that alter HIF TF activity

We confirmed 11 inhibitors and 18 activators of HIF1a activity identified from the pilot screen at three concentrations (**Supp Figure 8a**, **9a**) identifying RQ500235 and RQ200674 (**Figure 6a, d**) as previously unreported HIF-1α inhibiting or stabilising compounds, respectively. RQ200674 increased reporter activity 2-fold in repeated assays (**Figure 6d**) and stabilised endogenously tagged HIF-1α at normoxia in HEK293T cells (**Supp Figure 8b**). Mechanistically, RQ200674 had weak iron chelation activity in an *in vitro* chelation assay (**Figure 6e**), suggesting it intersects with the HIF-1α pathway by sequestering iron similar to other reported HIF stabilisers. In the inhibitor compound dataset, Celastarol and Flavokawain B downregulated the reporter at several concentrations (**Supp Figure 9b, c**). Celastarol is a previously reported HIF-1α inhibitor^46–48^ and Flavokawain B is a member of the chalcone family which has previously exhibited anti-HIF-1α activity^49^. RQ500235 was identified as a HIF-1 inhibitor by mcdFLASH-HIF screening. Dose dependent inhibition of mcdFLASH-HIF (**Figure 6a**) correlated with a dose-dependent decrease in protein expression by immunoblot (**Figure 6C**). We observed significant (p=0.0139) downregulation of HIF-1α transcript levels (**Figure 6D**) and were unable to rescue HIF-1α protein loss with proteasomal inhibition (**Supp Figure 9d**), indicating RQ500235 was decreasing HIF-1α at the RNA level. More broadly however, the identification of these compounds by mcdFLASH-HIF in the bimodal set up demonstrates successful application of the dFLASH platform to small molecule discovery efforts for both gain and loss of TF function.

**Figure 6.**
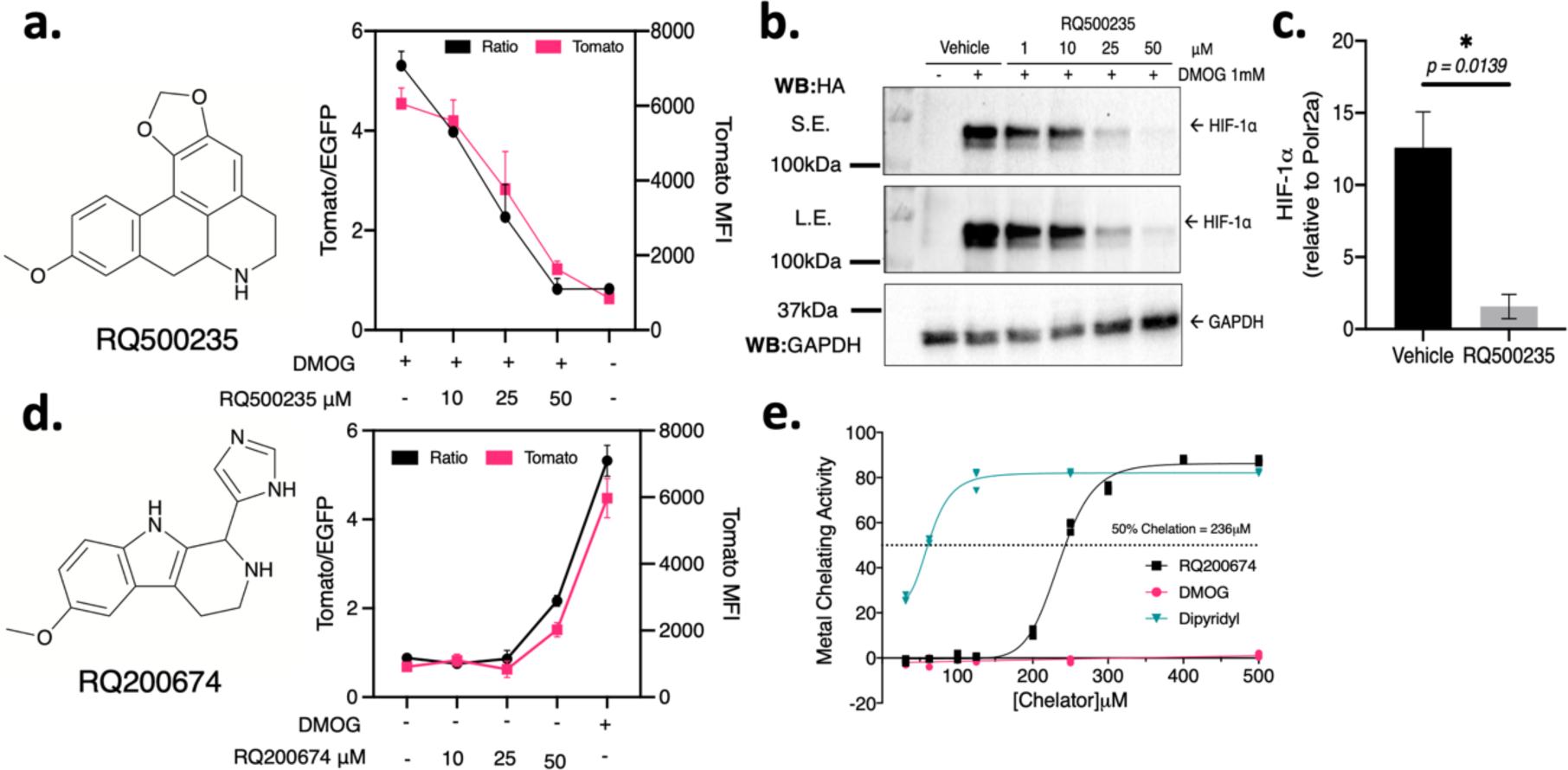
Investigating mechanisms for HIF-1α regulation by hit dFLASH-HIF inhibitor RQ500235 and hit activator RQ200674. (**a, b**) Inhibitor RQ500235 identified from the bimodal screen (**a**) represses DMOG induced Tomato in dFLASH-HIF cells in a dose dependent manner (n=2, Tom MFI, red; Tom normalised to EGFP, black) and (**b**) decreases expression of HIF-1α protein as assessed by immunoblot of whole cell extracts from endogenous HA-Flag tagged HIF-1α in HEK293T cells. S.E.= short exposure; L.E.= long exposure. (**c**) RT-PCR shows HIF-1α transcript is significantly decreased in HEK293T cells treated for 6 hours with RQ500235 (n =3, *p=0.0139). (**d**) Activator RQ200674 identified from the bimodal screen recapitulated activation of dFLASH-HIF at 50μM in HEK293T cells (n = 2). (**e**) *in vitro* iron chelation assay of RQ200674 displays weak chelating activity at 236μM from line of best fit (n = 3) compared to positive control iron chelator and HIF-1α activator, dipyridyl.

## Discussion

We designed and optimised dFLASH to offer a versatile, robust live-cell reporting platform that is applicable across TF families and allows for facile high-throughput applications. We validated dFLASH against three independent signal-responsive TFs, two with endogenous signalling pathways (dFLASH-PRE for Progesterone receptors; dFLASH-HRE for hypoxia induced transcription factors) and a synthetic system for a hybrid protein transcriptional regulator (dFLASH-Gal4RE). Each dFLASH construct produced robustly detected reporter activity by temporal high-content imaging and FACS after signal stimulation for its responsive TF (**Figure 2,3**). The use of previously validated enhancer elements for HIF^24^ and synthetic Gal4 DNA binding domains^22,28^ demonstrated that dFLASH can be adapted toward both endogenous and synthetic pathways displaying highly agonist/activator-specific responses, indicating utility in dissecting and targeting distinct molecular pathways. mcdFLASH lines distinct pathways produced highly consistent (Z’ = 0.68-0.74) signal induced Tomato induction measured by high content imaging suggesting dFLASH is ideally suited to arrayed high-throughput screening (**Figure 3**). In addition, mcdFLASH lines also displayed homogenous signal induced reporter induction by flow cytometry indicating that pooled high content screening would also be possible.

Indeed, reporter systems like dFLASH have been increasingly applied to functional genomic screens which target specific transcriptional pathways^9,50–52^. CRISPRoff mediated downregulation of the core HIF protein regulator, VHL produced distinct tomato expressing cell pools (**Figure 4**), demonstrating genetic perturbations of endogenous TF signalling pathways. The robust induction of the dFLASH-HIF reporter upon VHL knockdown in the majority of cells indicates that whole genome screening would also be successful^9,17,50,53^.

Using the HIF-1α specific reporter line, mcdFLASH-HIF, the application of high-content screening was exemplified. This approach was successful in discovering a novel activator and novel inhibitor of the HIF pathway, as well as previously identified inhibitory compounds. This ratified dFLASH as a reporter platform for arrayed-based screening and demonstrates the utility of the linked nucEGFP control for rapid hit bracketing. The novel inhibitor RQ500235 was shown to downregulate HIF-1α transcript levels, like another HIF-1α inhibitor PX-478^54,55^. As PX-478 has demonstrated anti-cancer activity in several cell lines ^55,56^ and preserved *β*-cell function in diabetic models^54^, a future similar role may exist for an optimised analogue of RQ500235.

The dFLASH system is characterised by some distinct advantages which may enable more precise dissection of molecular pathways. The ability to control for cell-to-cell fluctuations and to decouple generalised or off-target effects on reporter function may aid the precision necessary for large drug library or genome-wide screening applications^57^. In addition, dFLASH, unlike many other high-throughput platforms can be used to screen genetic or drug perturbations of temporal transcriptional dynamics or as used here at multiple time points. Also, the results indicate that dFLASH is ideally suited to array-based functional genomics approaches^58^ allowing for multiplexing with other phenotypic or molecular outputs^59,60^ ^2,61^.

The dFLASH approach has some limitations. The fluorescent nature of dFLASH limits the chemical space by which it can screen due to interference from auto-fluorescent compounds. In addition, we acknowledge that fluorescent proteins require O_2_ for their activity and this limits the use of mnucTomato as a readout of hypoxia. Also, while the backbone design has been optimised for a robust activation of a variety of transcription response pathways, the mechanistic underpinning of this is unclear and could be further improved, providing insights into the sequence and architectural determinants of enhancer activation in chromatin. In addition to the strong effect of the dFLASH downstream promoter on upstream enhancer activity it is clear that either the distance between contiguous promoter/enhancer or the sequence composition of the linker has a functional consequence on enhancer induction.

The incorporation of robust native circuits such as those described here (Hypoxia or Progesterone) has the potential to allow the manipulation or integration of these pathways into synthetic biology circuitry for biotherapeutics. In these cases, it is critical that robust signal to noise is achieved for these circuits to effectively function in biological systems. Further, the use of a synthetic approach to ‘sense’ FIH enzymatic activity through the HIF-CAD:P300/CBP interaction opens up the possibility that other enzymatic pathways that lack effective *in vivo* activity assay may also be adapted. We also envisage that dFLASH could be adapted to 2-hybrid based screens as a complement to other protein-protein interaction approaches.

The ability to temporally track TF regulated reporters in populations and at the single-cell level enable dFLASH to be used to understand dynamics of transcriptional responses as has been used to dissect mechanisms of synthetic transcriptional repression^7,8^ or understand notch ligand induced synthetic transcriptional dynamics^62^. For instance, synthetic reporter circuits have been used to delineate how diverse notch ligands induce different signalling dynamics ^62^. The large dynamic range of the dFLASH-PGR and HIF reporter lines in conjunction with the high proportion of cells induced in polyclonal pools (**Figure 2**) also suggests dFLASH as a candidate system for forward activity-based enhancer screening. These approaches have been applied to dissect enhancer activity or disease variants with other similar systems such as lentiviral-compatible Massively Parallel Reporter Assays (LentiMPRA)^63,64^. However, the use of the internal control normalisation provided by dFLASH may be useful in separating chromosomal from enhancer driven effects in forward enhancer screens.

Given dFLASH has robust activity in both pooled and arrayed formats, it offers a flexible platform for investigations. dFLASH can be used to sense endogenous and synthetic transcription factor activity and represents a versatile, stable, live-cell reporter system of a broad range of applications.

## Supplementary Figures

**Supplementary Figure 1.**
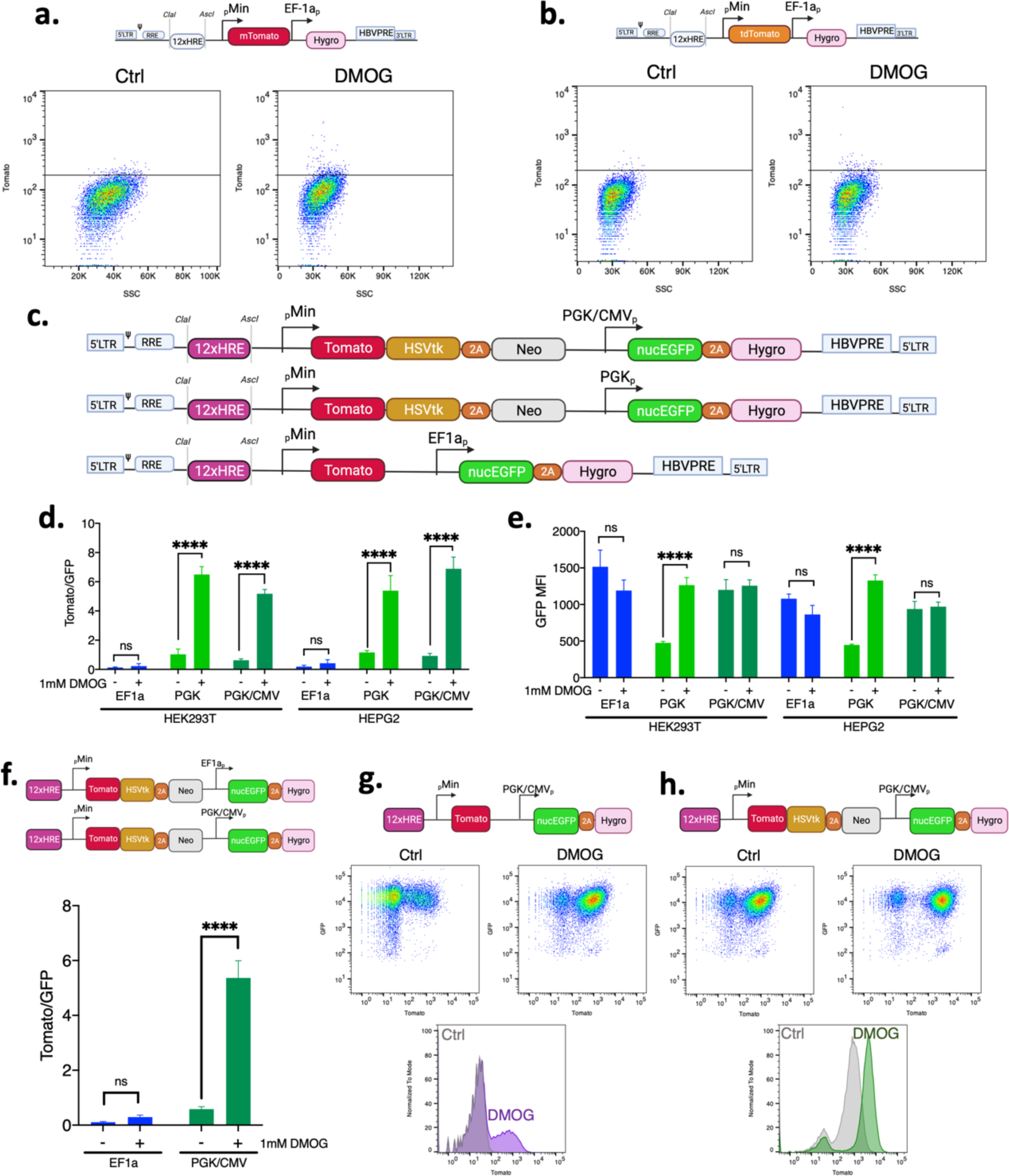
Optimised dFLASH design produces a robust HIF sensor. (**a-b**) HEK293T cells with HRE-dFLASH constructs without EGFP and (**a**) expressing monomeric Tomato or (**b**) dimeric Tomato were treated −/+ 1mM DMOG for 48 hours and quantified by FACS. Tomato MFI >200AU was used to compare induction (black line). (**c-e**) HEK293T and HEPG2 cells were transduced with HRE-dFLASH reporters that had different downstream promoters controlling EGFP or Tomato cassette composition and treated for 48 hours −/+ 1mM DMOG prior to HCI. The (**d**) Tomato/EGFP MFI ratio and (**e**) EGFP MFI for each backbone variant was then compared (Data from three independent biological replicates). (**f**) HEK293T cells transduced with reporter constructs containing the downstream PGK/CMV or EF1a promoters were compared for DMOG induction by HCI after 48 hours of −/+ 1mM DMOG treatment (Data from three independent biological replicates). Significance was assessed with a Two-Way ANOVA (**** p < 0.001, ns = not significant). (**g,h**) HEK293T cells with the HRE enhancer and different dFLASH backbone compositions of (**g**) PGK/CMV dFLASH with Tomato alone as the upstream cassette or (**h**) dFLASH-HIF were treated for 48-hours −/+ 1mM DMOG prior to EGFP analysis and Tomato induction by FACS.

**Supplementary Figure 2.**
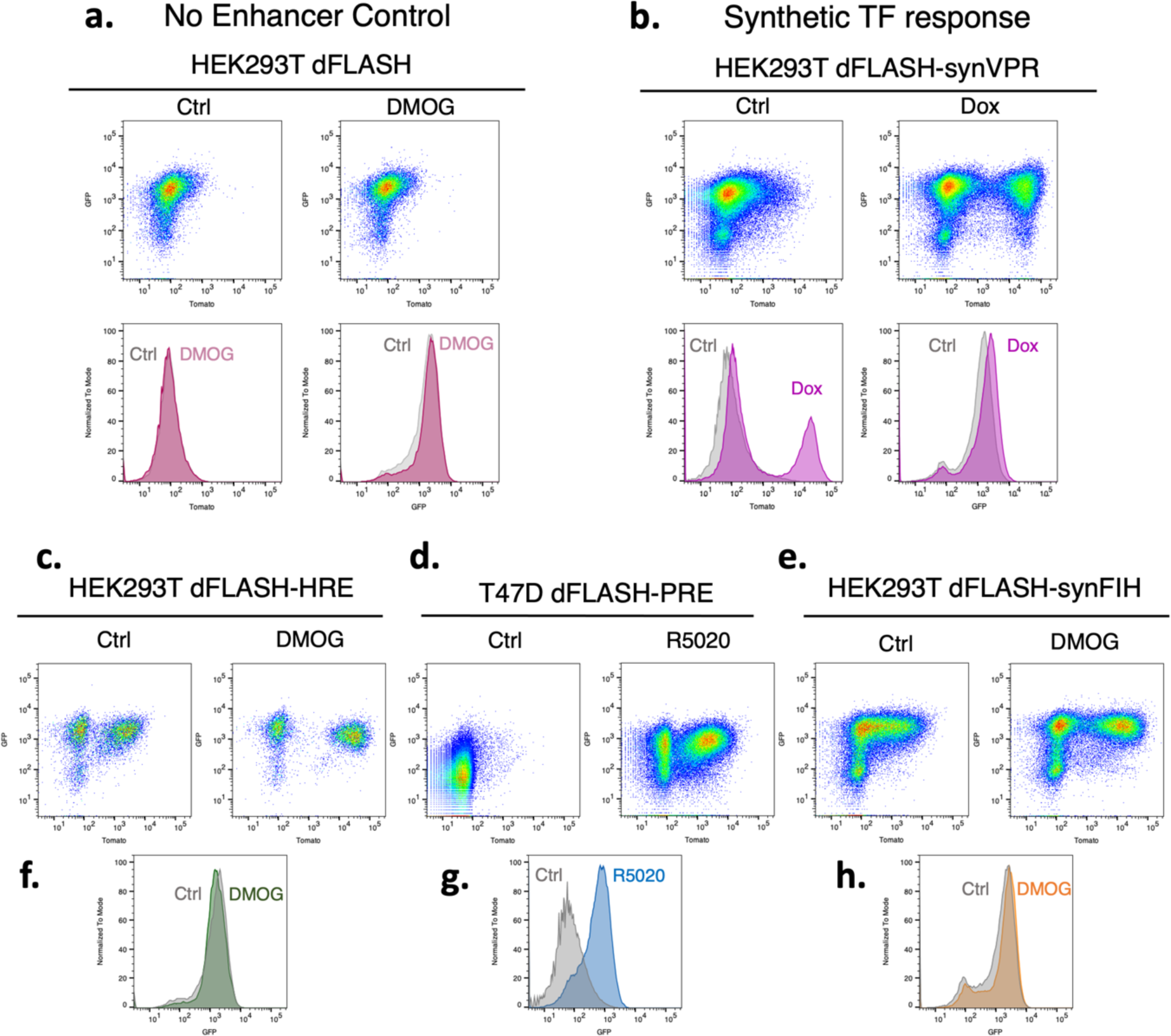
dFLASH provides a TF-responsive, versatile reporter platform in heterogenous cell pools. (**a-b**) HEK293T cells were transduced with (**a**) dFLASH with no enhancer and treated with 1mM DMOG or 0.1% DMSO (Ctrl) or (**b**) GalRE-dFLASH and Gal4DBD-miniVPR and treated with H_2_O (Ctrl) or 1μg/mL Dox for 48 hours prior to FACS. Dot plots of populations’ Tomato and EGFP intensity with or without activating chemicals and histograms comparing EGFP and Tomato MFI between control and treated populations are shown. (**c-h**) Dot plots and EGFP histograms for control and chemical treated (**c, f**) dFLASH-HIF, (**d, g**) dFLASH-PR polyclonal pools (*to accompany* Figure 2a-c) and (**e, h**) dFLASH-synFIH.

**Supplementary Figure 3.**
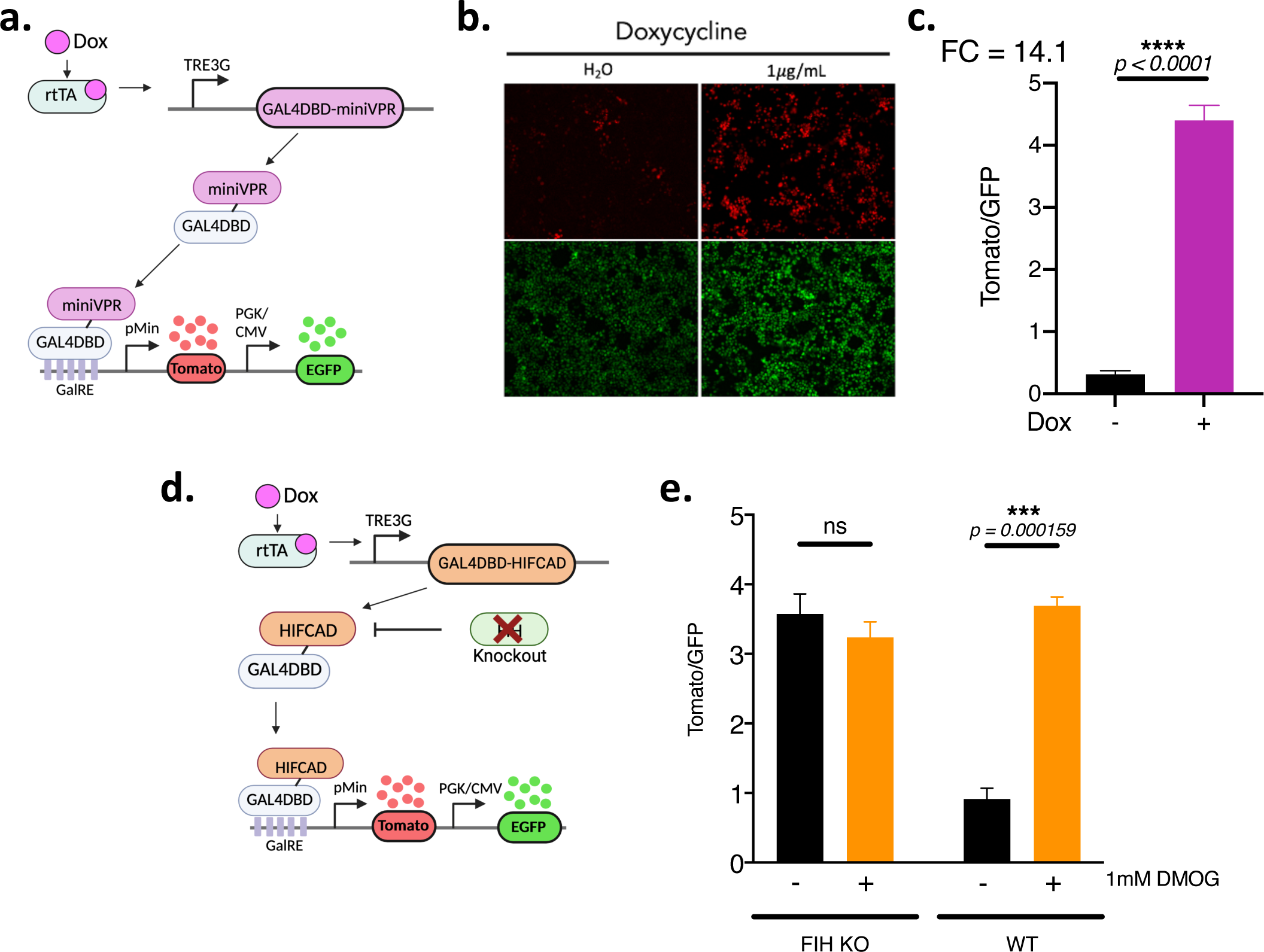
Synthetic transcription factors drive a strong response from the GalRE-dFLASH reporter and can respond to endogenous signaling pathways. (**a**) GAL4DBD-miniVPR is expressed from an independent dox-inducible vector that subsequently binds to GalRE-dFLASH. (**b**,**c**) HEK293T GalRE-dFLASH cells were transduced with GAL4DBD-miniVPR expression construct and were treated −/+ doxycycline for 48 hours prior to HCI for (**b**) Tomato expression (top panels) and EGFP expression (bottom panels). (**c**) Normalised fluorescence intensity was also quantified for treated populations (n=3, mean ±sem). FC is Fold change between the populations. (**d, e**) To confirm HEK293T dFLASH-synFIH system was FIH dependent, (**d**) GalRE-dFLASH and GAL4DBD-HIFCAD vectors were transduced into HEK293T cells with FIH knocked out. (**e**) FIH KO cells were compared with wildtype HEK293T dFLASH-synFIH (WT) in a 200ng/mL dox background for DMOG-dependent reporter induction by HCI (n=3). (**c, e**) Significance was assessed by t-test with Welch’s correction (ns = not significant, *** p <0.001, ****p <0.0001).

**Supplementary Figure 4.**
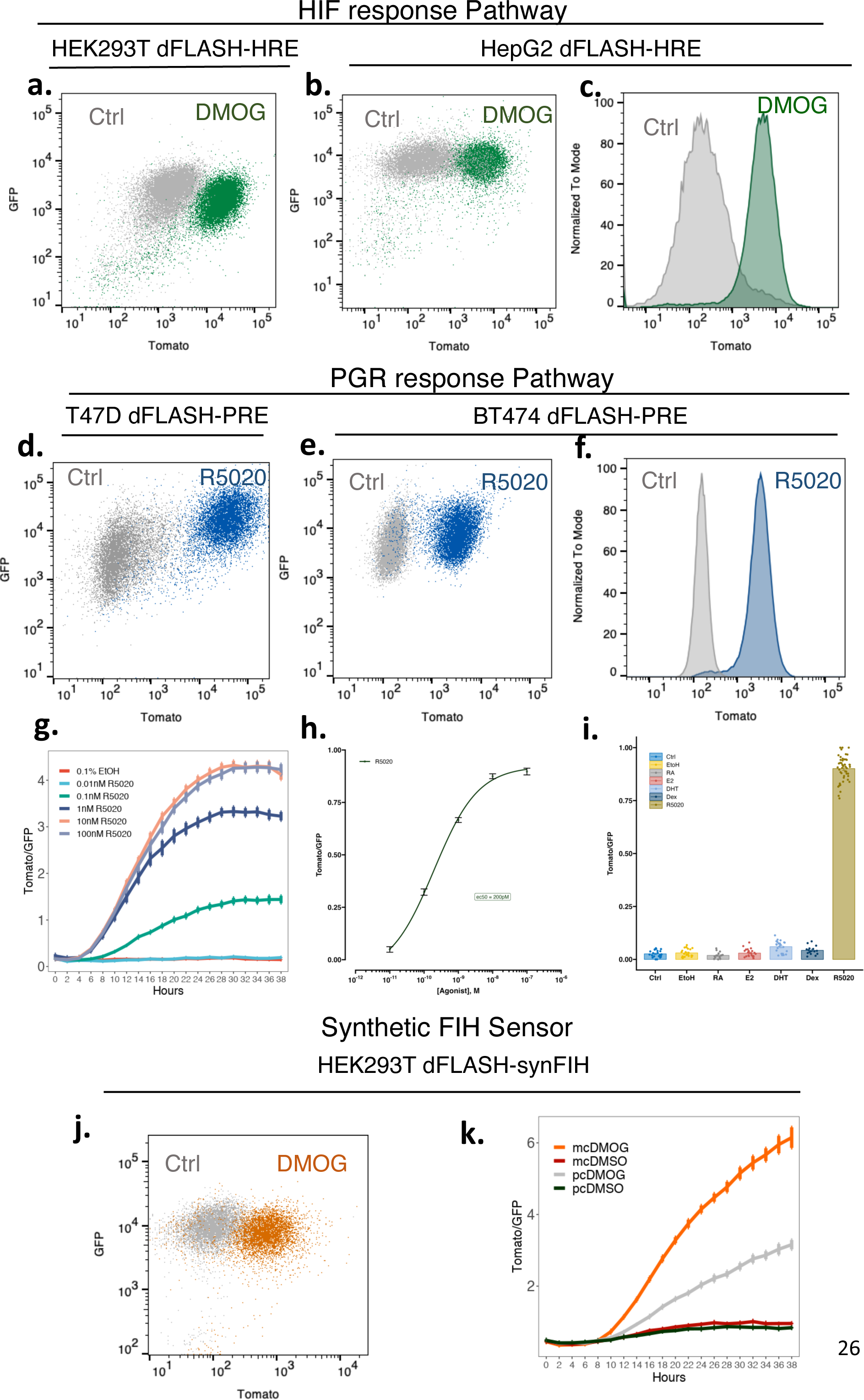
Clonal dFLASH cell lines enable improved reporting across different cell types. (**a-c**) Flow cytometry of clonal dFLASH-HIF cell lines for (**a**) HEK293T (*see also* Figure 3b) and (**b,c**) HepG2 cells after 48 hours −/+ 0.5mM DMOG. (**d-h**) dFLASH-PGR functionality was assessed by flow cytometry in (**d**)T47D (*see also* Figure 3f) and (**e,f**) BT474 cells after 48 hours −/+ 100nM R5020. (**g,h**) T47D dFLASH-PGR cells were treated with increasing concentrations of R5020 (0.01-100nM, 8 replicates per group) and (**g**) imaged over 38 hours with temporal HCI or (**h**) imaged at 48 hours to determine sensitivity to R5020. (**i**) Comparison of inductions of the T47D mcdFLASH-PGR line to different steroids (10nM R5020, 35nM E2, 10nM DHT, 10nM Dex, 10nM RA) by HCI after 48 hours of treatment. (**g**) and (**i**) are the mean±sem of normalised Tomato/GFP (within each expt) from n = 3 independent experiments (24 replicates), except Dex and RA (n=2 (16 replicates)). (**j, k**) Clonally derived HEK293T dFLASH-synFIH cells were (**j**) analysed by flow cytometry after 48 hours of 200ng/mL Dox −/+ 1mM DMOG (*see also* Figure 3k) with (**k**) showing temporal HCI comparisons between monoclonal (mc) and polyclonal (pc) lines (*see also* Figure 2j).

**Supplementary Figure 5.**
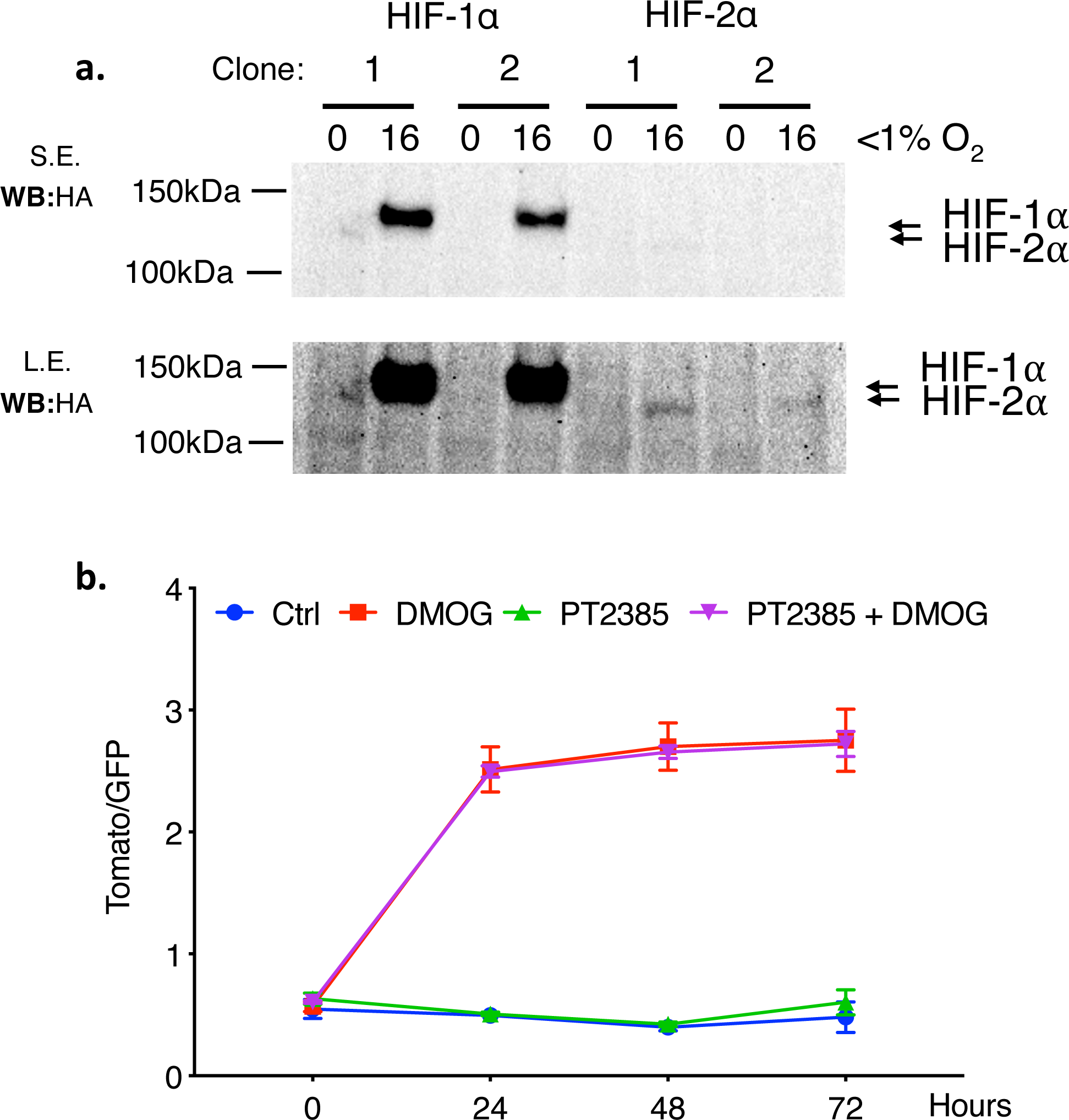
HIF-1α is the predominant isoform that affects the dFLASH reporter in HEK293T cells. (**a**) Monoclonal HEK293T cells with endogenously HA-Flag tagged HIF-1α or HIF-2α were treated with hypoxia (<1% O_2_) for 16 hours prior to anti-HA immunoblotting of whole cell extracts. S.E.= short exposure; L.E.= long exposure. Representative of three independent experiments. (**b**) mcdFLASH-HIF cells were treated −/+ 1mM DMOG and −/+ 10μM of the HIF-2α antagonist (PT-2385) as indicated and quantified by HCI over 72-hour period.

**Supplementary Figure 6.**
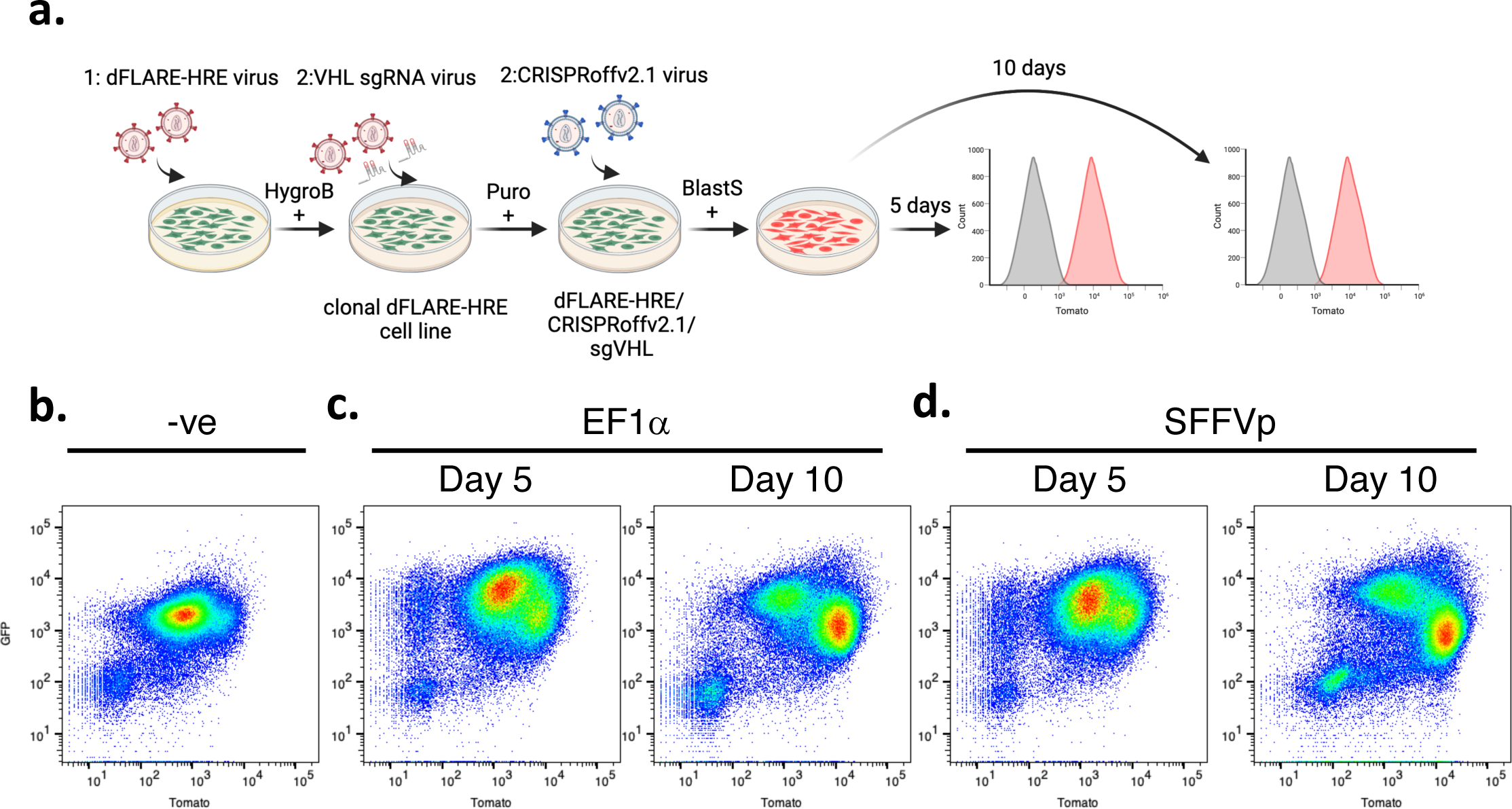
CRISPRoff mediated VHL knockdown induces mcdFLASH-HIF reporter lines. (**a**) HEK293T cells were first transduced with dFLASH-HRE and a clonal reporting line was derived after hygromycin (HygroB) treatment. This line was in turn transduced with the VHL sgRNA vector and selected with puromycin (Puro). This line was then transduced with the CRISPRoffv2.1 vector and selected with blasticidin S (Blast) and populations were subjected to flow cytometry after 5 days or 10 days of selection for analysis of reporter expression. (**b-d**) dot plots for dFLASH expression from the (**b**) non-CRISPRoff parental line, (**c**) EF1a-CRISPRoffv2.1 transduced and (**d**) SFFVp-CRISPRoffv2.1 populations after 5 or 10 days of blasticidin selection (*see also* Figure 4).

**Supplementary Figure 7.**
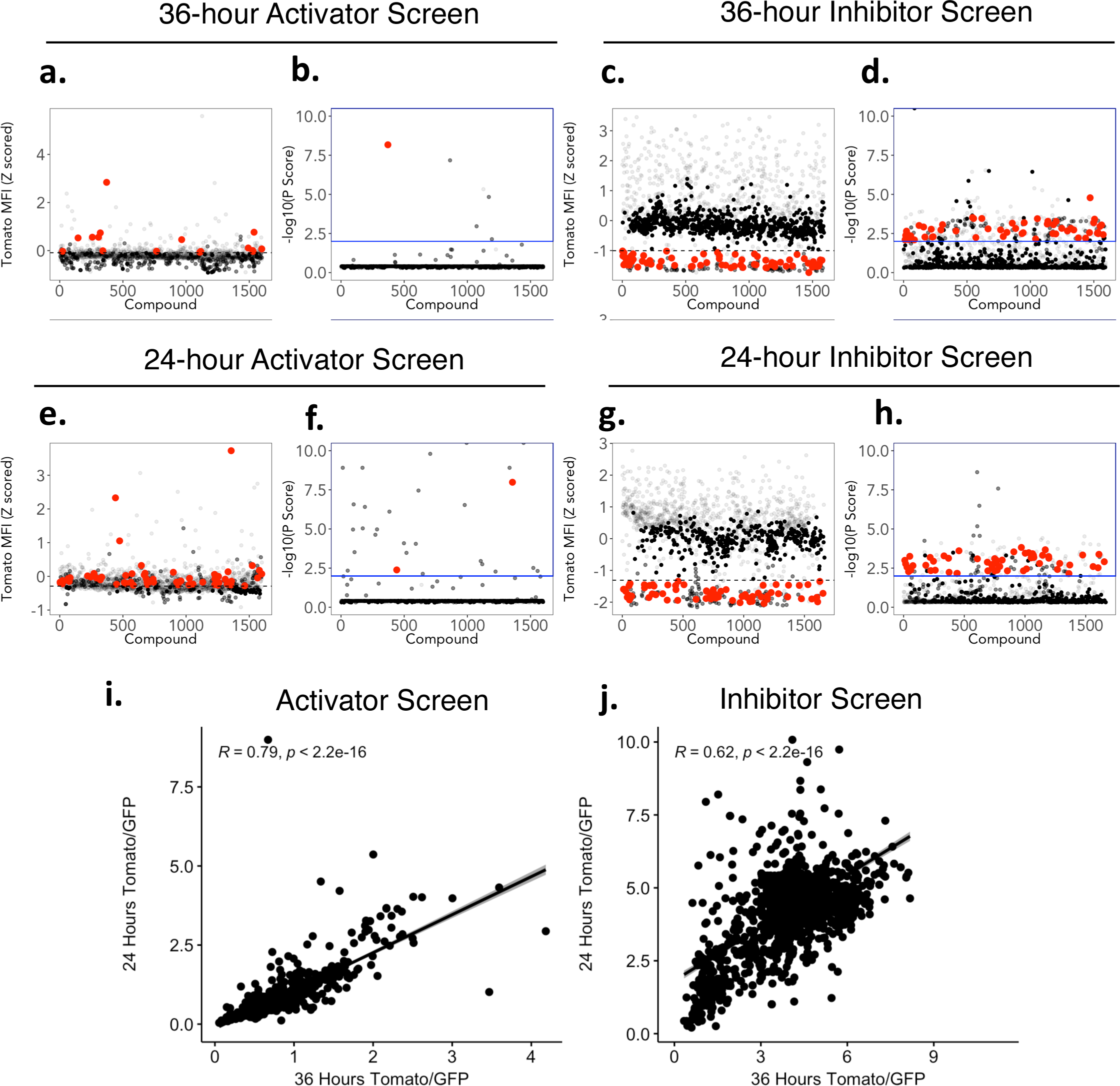
Hit selections and assessment of bimodal screen reproducibility between independent screens for activators and inhibitors of HIF-1α. Compound-induced dFLASH-HIF reporter activity was used to score hits from the (**a-d**) 36-hour or the (**e-h**) 24-hour bimodal screens according to Tomato MFI and adjusted P scores. Lines indicate cut offs for hit criteria with hits shown in red for each metric and dismissed compounds that change EGFP >±2SD shown in grey. (**i, j**) Pearson correlations of the Tomato/EGFP between the 36-hour and the 24-hour screens for (**i**) reporter activation (R = 0.62, p < 2.2×10^−16^) or (**j**) reporter inhibition (R = 0.62, p < 2.2×10^−16^) for all 1595 compounds screened. Line indicates line of best fit, grey boundary is 95% confidence interval.

**Supplementary Figure 8.**
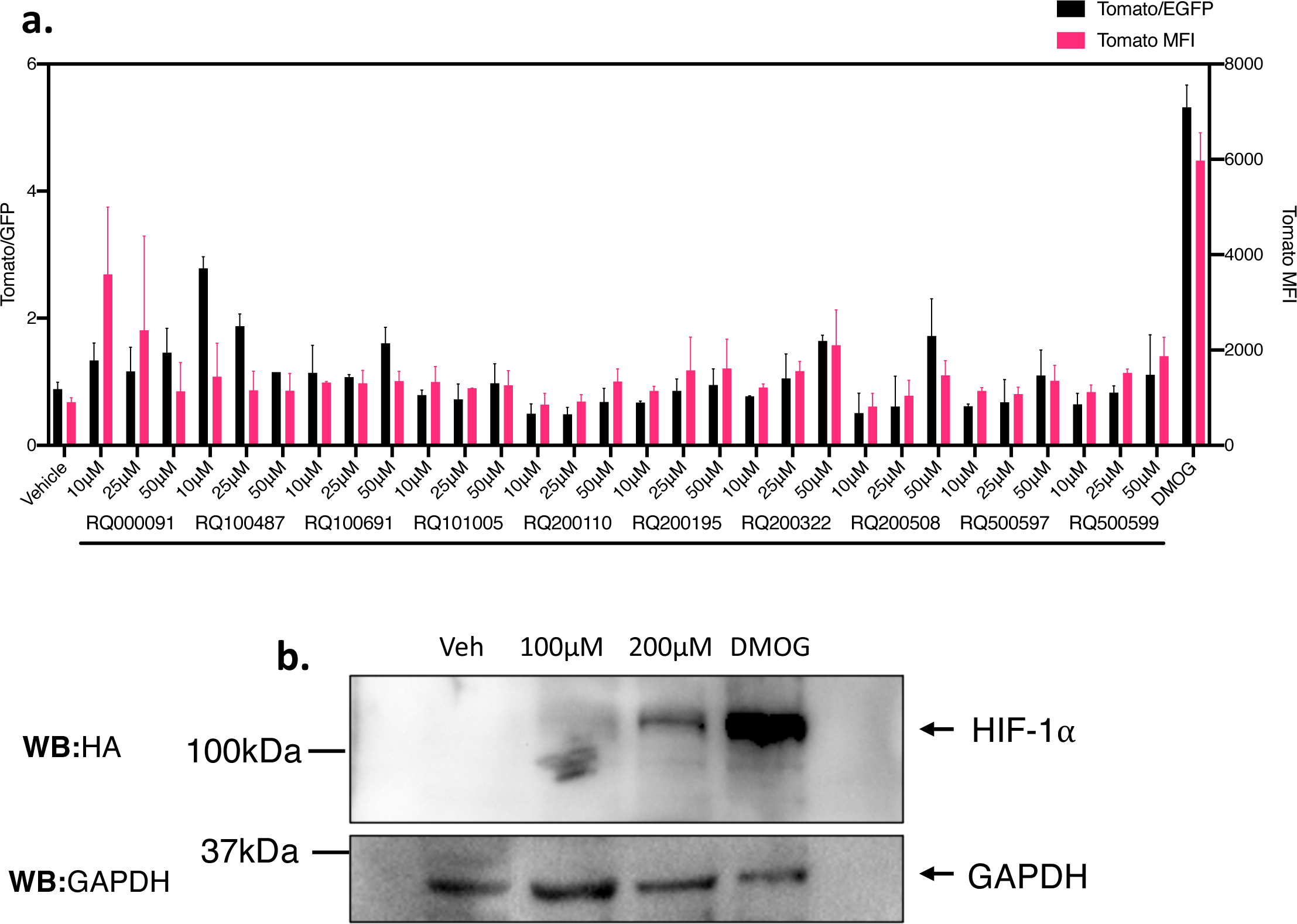
Rescreening of activator hits from 1595 compound small molecule screen reveals RQ200674 causes normoxic stabilisation of HIF-1α. **(a)** The 11 top performing hits from the activator screens, including RQ200674 (*see also* Figure 6d) were rescreened against HEK293T mcdFLASH-HIF at 10*μ*M, 25*μ*M and 50*μ*M. Comparisons between Tomato/GFP and Tomato MFI dFLASH induction shown against vehicle (-ve Ctrl) and 1mM DMOG (+ve Ctrl) treated populations (n=2). **(b)** Immunoblot of whole cell extracts from HEK293T cells containing endogenously HA-Flag tagged HIF-1α and treated as indicated with vehicle (0.1% DMSO),1mM DMOG (+ve Ctrl), or 100*μ*M and 200*μ*M of RQ200674 for 18 hours. Representative of 2 independent experiments.

**Supplementary Figure 9.**
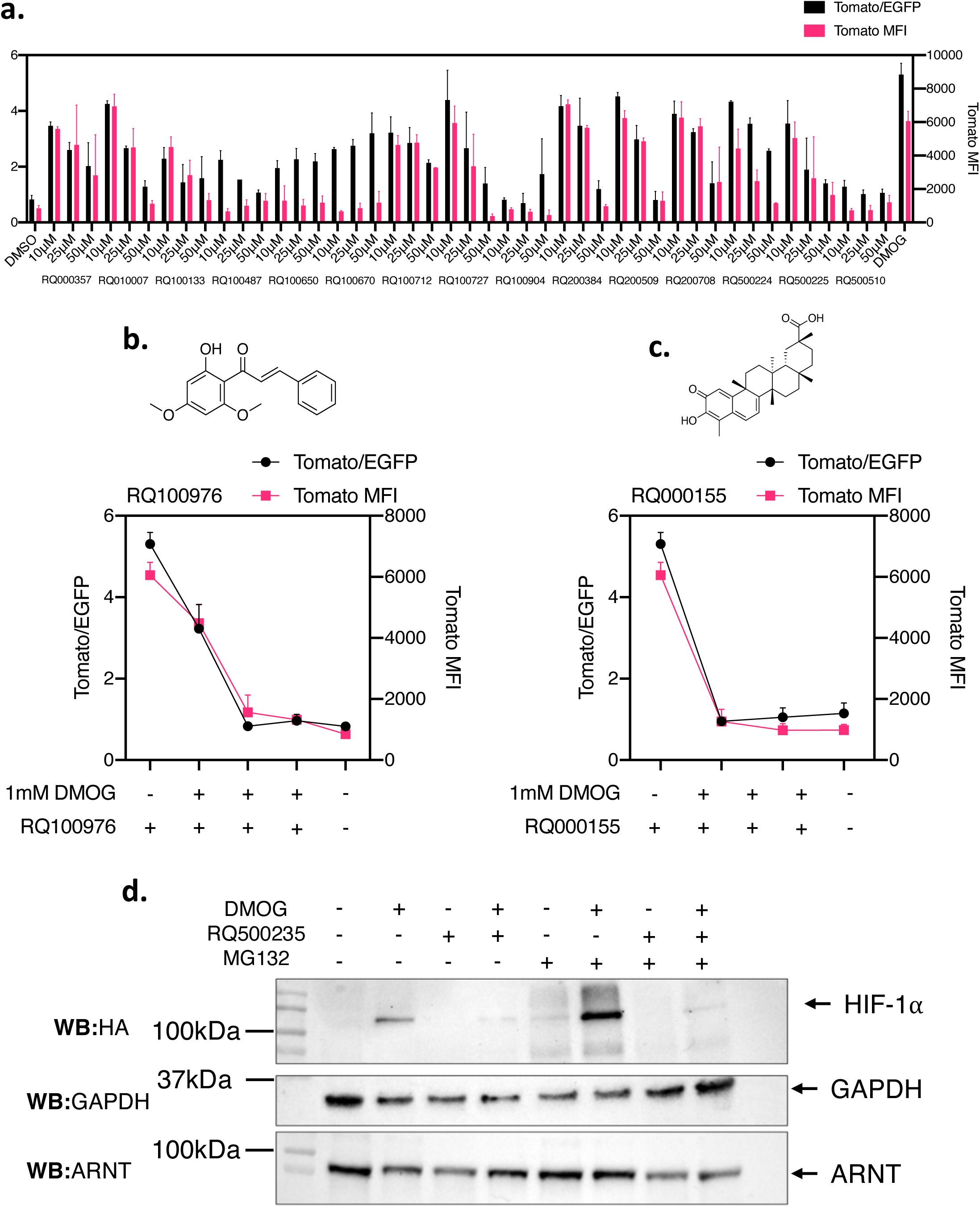
Flavokawain B, Celastarol and RQ500235 decrease dFLASH-HIF and proteasomal inhibition doesn’t rescue RQ500235 impact on HIF-1α. (**a-c**) The 18 top inhibitory compounds, including (**b**) Flavokawain B (RQ100976),(**c**) Celastarol (RQ000155) and RQ500235 (*see also* Figure 6a) were rescreened against dFLASH-HIF at 10μM, 25μM and 50μM in 1mM DMOG treated 293T dFLASH-HIF cells (24 hours). Comparisons between Tomato/GFP and Tomato MFI dFLASH induction shown against 0.1% DMSO (-ve Ctrl) and 1mM DMOG (+ve Ctrl) treated populations (n=2). (**d**) Immunoblot of whole cell extracts from HEK293T cells with endogenously HA-Flag tagged HIF-1α following a 12 hr treatment period with with the indicated combinations of 1 mM DMOG (full12 hr), 50*μ*M RQ500235 (final 6 hr) and 10*μ*M MG132 (final 3 hr). Representative of 2 independent experiments.

**Supplementary Movie 1.**
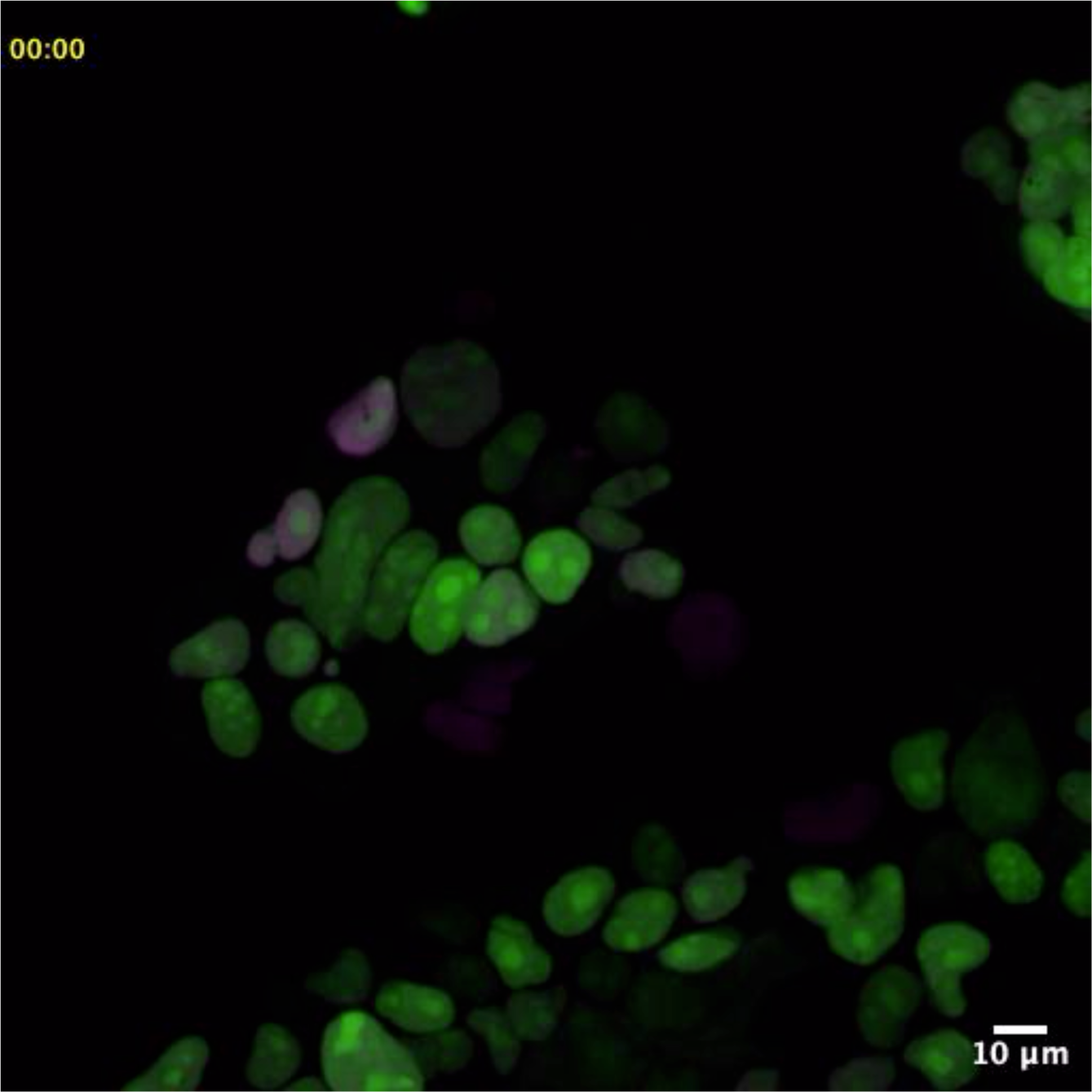
Single cell temporal dynamics of HEK293T mcdFLASH-HIF cells. HEK293T mcdFLASH-HIF cells were seeded at 1×10^5^ cells/dish in Poly-D-Lysine coated plates overnight prior to imaging with spinning disk confocal microscopy at 40x magnification. Cells were imaged every 15 min for 48 hours for Tomato (Magenta) and EGFP (Green) expression. Time stamps are given in top left.

**Supplementary Movie 2.**
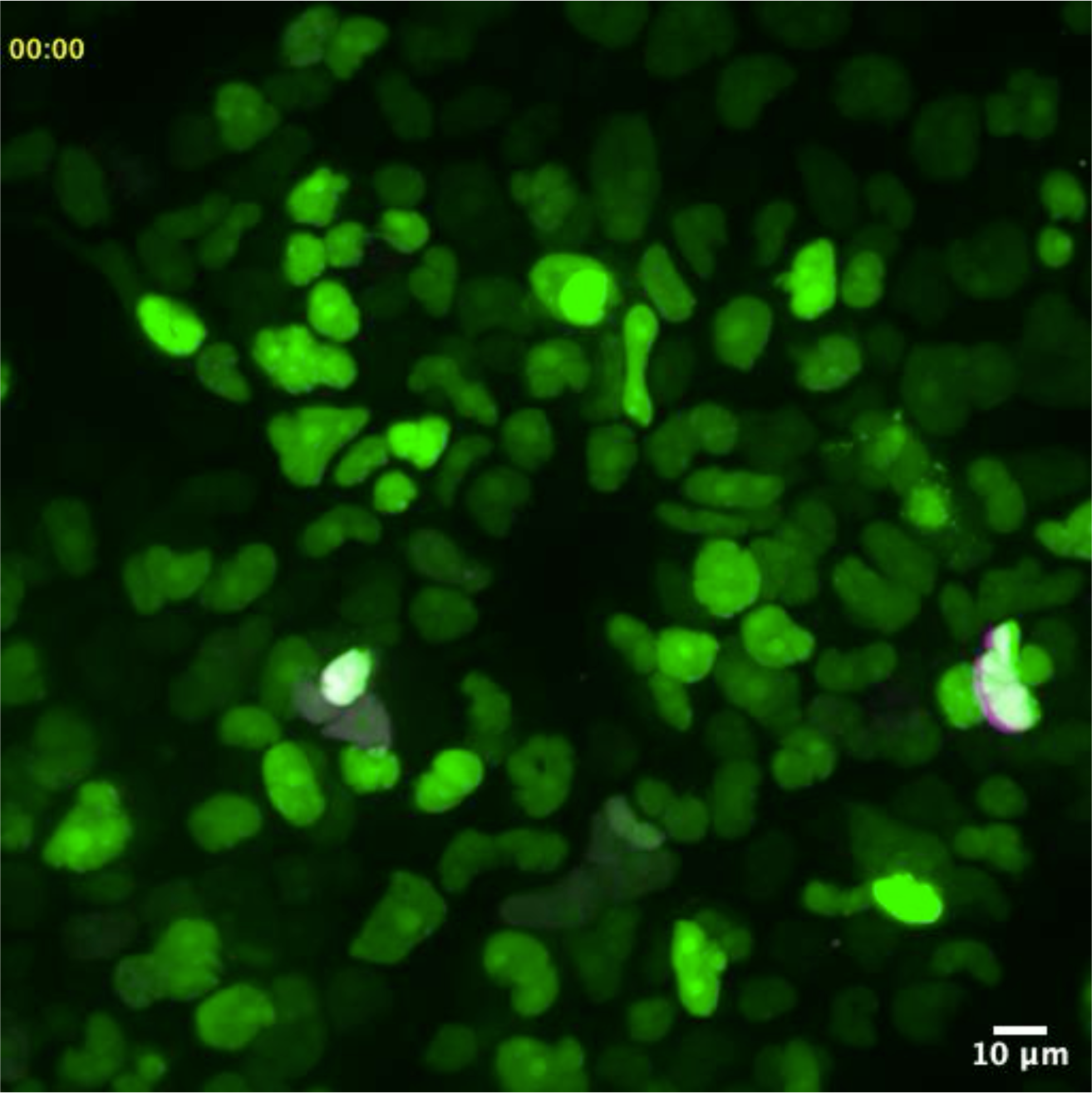
Single cell temporal dynamics of T47D mcdFLASH-PGR cells. T47D mcdFLASH-PGR cells were seeded at 5×10^5^ cells/dish in Poly-D-Lysine coated plates overnight prior to imaging with spinning disk confocal microscopy at 40x magnification. Cells were imaged every 15 min for 48 hours for Tomato (Magenta) and EGFP (Green) expression. Time stamps are given in top left.

## Methods

### Plasmid Construction

cDNAs were amplified using the Phusion polymerase (NEB) and assembled into ClaI/NheI digested pLV410 digested backbone by Gibson assembly^31^. Sequence verified LV-REPORT plasmid sequences and constructs are listed in **Supplementary Table 1**. Briefly, the plasmids contained an upstream multiple cloning sites followed by a minimal promoter (derived from the pTRE3G minimal promoter) and then followed by a reporter construct mnucTomato/HSVtk-2a-Neo or other variants). This was then followed by a constitutive promoter (EF1a, PGK or PGK/CMV) driving the expression or hygromycinR cassette with or without a 2a linked d2nucEGFP (**Supplementary Figure 1C**).

To improve the performance of our previously reported lentiviral inducible expression systems^65^, the PGK promoter in Tet-On3G IRES Puro was replaced by digestion with MluI/NheI and insertion of either EF1a-Tet-On3G-2A-puro, EF1a-Tet-On3G-2A-BlastR or EF1a-Tet-On3G-2A-nucTomato using Phusion polymerase (NEB) amplified PCR products from existing plasmids. Plasmids were cloned by Gibson isothermal assembly and propagated in DB3.1 cells (Invitrogen). We also generated a series of constitutive lentiviral plasmids as part of this work pLV-EgI-BlastR (EF1a-Gateway-IRES-BlastR), pLV-EgI-ZeoR (EF1a-Gateway-IRES-ZeoR), pLV-EgI-HygroR (EF1a-Gateway-IRES-HygroR), pLV-SFFVp-gI-BlastR (SFFVp-Gateway-IRES-BlastR), pLV-SV40p-gI-BlastR (SV40p-Gateway-IRES-BlastR). These plasmids were constructed by isothermal assembly of G-Blocks (IDT DNA) or PCR fragments, propagated in ccbD competent cells, sequence verified and deposited with Addgene (**Supplementary Table 1**).

The Lentiviral backbone expression construct pLV-TET2BLAST-GtwyA was then using to insert expression constructs cloned into pENTR1a by LR Clonase II enzyme recombination (Cat#11791020, Thermo). GAL4DBD-HIFCAD (727-826aa) and the GAL4DBD^28^ were cloned into pENTR1a by ScaI/EcoRV or KpnI/EcoRI respectively. The miniVPR sequence^29^ was cloned into the pENTR1a-GAL4DBD construct at the EcoRI and NotI sites. The pENTR1a vectors were then Gateway cloned into the pLV-TET2PURO-GtwyA vector. pENTR1a-CRISPRoffv2.1 was generated by inserting an EcoRI/NotI digested CRISPRoff2.1 (CRISPRoff-v2.1 was a gift from Luke Gilbert, Addgene #167981) into pENTR1a plasmid. pLV-SFFVp-CRISPRofv2.1-IRES-BLAST and pLV-EF1a-CRISPRofv2.1-IRES-BLAST were generated by pENTR1a by LR Clonase II enzyme recombination (Cat#11791020, Thermo). All Lentiviral plasmids were propagated in DH5a without any signs of recombination.

### Enhancer element cloning

The 12x HRE enhancer from hypoxic response target genes (PGK1, ENO1 and LDHA) was liberated from pUSTdS-HRE12-mCMV-lacZ^24^ with XbaI/SpeI and cloned into AvrII digested pLV-REPORT plasmids. Progesterone responsive pLV-REPORT-PRECat PRECat was cloned by isothermal assembly of a G-Block (IDT-DNA) containing enhancer elements from 5 PGR target gene enhancers (Zbtb16, Fkbp5, Slc17a11, Erfnb1, MT2)^66^ into AscI/ClaI digested pLV-REPORT(PGK/CMV). Gal4 response elements (5xGRE) were synthesised (IDT DNA) with ClaI/AscI overhangs and cloned into Cla/AscI digested pLV-REPORT(PGK/CMV). Sequences are in **Supplementary Table 2.**

### Mammalian cell culture and ligand treatment

HEK293T (ATCC CRL-3216), HEPG2 (ATCC HB-8065) line were grown in Dulbecco’s Modified Eagle Medium (DMEM high glucose) + pH 7.5 HEPES (Gibco), 10% Foetal Bovine Serum (Corning 35-076-CV or Serana FBS-AU-015), 1% penicillin-streptomycin (Invitrogen) and 1% Glutamax (Gibco). T47D (ATCC HTB-133) or BT474 (ATCC HTB-20) were grown in RPMI 1640 (ATCC modified) (A1049101 Gibco) with 10% Foetal Bovine Serum (Fisher Biotech FBS-AU-015) and 1% penicillin-streptomycin^67^. Cells were maintained at 37^0^C and at 5% CO_2_.Clonal lines were isolated by either limiting dilution or FACS single cell isolation into 96 wells trays. Resultant monoclonal populations were evaluated for single colony formation or assessed by HCI or FACS. Ligand treatments were done 24 hours after seeding of cells in requisite plate or vessel. Standard concentrations and solvent, unless specified otherwise, are 200ng/mL Doxycycline (Sigma, H_2_O), 0.5mM or 1mM DMOG (Cayman Scientific, DMSO), 100nM R5020 (Perkin-Elmer NLP004005MG, EtoH), 35nM Estradiol (E2, Sigma E2758, EtOH), 10nM all-trans retinoic acid (RA, Sigma #R2625), 10nM Dihydrotestosterone (DHT, D5027), 10nM Dexamethasone (Dex, Sigma D4902), 10µM PT-2385 (Abcam, DMSO).

### Lentiviral Production & stable cell line production

Near confluent HEK293T cells were transfected with either psPAX2 (Addgene #12260) and pMD2.G (Addgene #12259) or pCMV-dR8.2 dvpr (Addgene #8455), pRSV-REV (Addgene; #12253) and pMD2.G along with the Lentivector (described above) and PEI (1µg/µl, polyethyleneimine) (Polysciences, USA), Lipofectamine 2000, or Lipofectamine 3000 at a 3µl:1µg ratio with DNA. Media changed 1-day post-transfection to complete media or Optimem. Virus was harvested 1-2 days post-transfection, then viral media was filtered (0.45μM or 0.22μM, Sartorius) before the target cell population was transduced at a MOI < 1. Cells were incubated with virus for 48 hours prior media being exchanged for antibiotic containing complete media. Standard antibiotic concentrations were 140µg/mL hygromycin (ThermoFisher Scientific #10687010), 1µg/mL Puromycin (Sigma; #P8833) or 10µg/mL Blasticidin S (Sigma; CAT#15205).

### Generation of CRISPR knockout or knockdown cell lines

Generation of CRISPR knockout guides and plasmids against FIH has been previously described^68^. These guides were transfected into HEK293T cells and with PEI at a 3μg:1μg ratio then clonally isolated as above. Knockouts were confirmed with PCR amplification and sanger sequencing coupled with CRISPR-ID^69^. FIH knockouts were selected via serial dilution and confirmation of knockout by sequencing and T7E1 assay. The VHL sgRNA guides were selected from the Dolcetto CRISPRi library^70^ with BsmBI compatible overhangs (**Supplementary Table 3**). These oligos were annealed, phosphorylated then ligated into BsmBI-digested pXPR050 (Addgene#9692), generating XPR-050-VHL. Monoclonal HEK293T LV-REPORT-12xHRE cell lines were transduced with the XPR-050-sgVHL virus, and stable cell lines selected with Puromycin. Subsequently, LV-SFFVp-CRISPRoffv2.1-IRES-BlastR or LV-EF1a-CRISPRoffv2.1-IRES-BlastR virus was infected into HEK293T LV-REPORT-12xHRE/XPR-050-sgVHL stable cells and selected with Blasticidin S (15μg/ml) for 5 days. FACS was used to assess activation of the dFLASH-HRE reporter in parental (dFLASH-HRE/sgVHL) or CRISPRoffv2.1 expressing cells at day 5 or day 10 after Blasticidin S addition.

### CRISPR knock-in of tags to endogenous HIF-1α and HIF-2α

CRISPR targeting constructs clones targeting adjacent to the endogenous HIF-1α and HIF-2α stop codons^71^. Constructs were cloned into px330 by ligating annealed and phosphorylated oligos with BbsI digested px330, using hHIF-1α and hHIF-2α CTD sgRNA (**Supplementary Table 3**). Knock-in of HA-3xFlag epitopes into the endogenous HIF-1α or HIF-2α loci in HEK293T cells was achieved by transfection with 0.625 μg of pNSEN, 0.625μg of pEFIRES-puro6, 2.5μg of px330-sgHIF-α CTD, and 1.25μg of ssDNA HDR template oligo containing flanking homology to CRISPR targeting site the tag insertion and a PAM mutant into ∼0.8×10^6^ cells using PEI (3:1). 48 hours after transfection, the medium was removed from cells and replaced with fresh medium supplemented with 2 μg/ml puromycin for 48 hours and the cell medium was changed to fresh medium without puromycin. 48 hours later cells were seeded by limiting dilution into 96-well plates at an average of 0.5 cells/well. Correct integration was identified by PCR screening using HIF-1α and HIF-2α gDNA screening primers (**Supplementary Table 4).** Positive colonies reisolated as single colonies by limiting dilution. Isolated HIF-1α and HIF-2α tag insertions were confirmed by PCR, sanger sequencing and western blotting.

### High Content Imaging (HCI)

Cells were routinely seeded at 1×10^4^ to 5×10^4^ cells per well in black walled clear bottom 96 well plates (Costar Cat#3603), unless otherwise stated. Cell populations were imaged in media at the designated time points at 10x magnification and 2×2 binning using the ArrayScan^TM^ XTI High Content Reader (ThermoFisher). Tomato MFI and EGFP MFI was imaged with an excitation source of 560/25nm and 485/20nm respectively. Individual nuclei were defined by nuclear EGFP expression, nuclear segmentation and confirmed to be single cells by isodata thresholding. Nuclei were excluded from analysis when they couldn’t be accurately separated from neighbouring cells and background objects, cells on image edges and abnormal nuclei were also excluded. EGFP and Tomato intensity was then measured for each individual nucleus from at least 2000 individual nuclei per well. Fixed exposure times were selected based on 10-35% peak target range. Quantification of the images utilised HCS Studio^TM^ 3.0 Cell Analysis Software (ThermoFisher). For assessment of high throughput robustness of each individual reporting line in a high throughput setting (HTS-HCI), replicate 96 well plates were seeded for the HIF (10 plates), PGR (5 plates) and synFIH (3 plates) monoclonal reporter lines and imaged as above at 48 hours. For the HIF line, each plate had 6 replicates per treatment (vehicle or DMOG) per plate. For the PGR, 24 replicates per treatment, either vehicle or R5020 per plate were present with edge wells excluded. 24 replicates per treatment were also used for synFIH, with system robustness assessed between the DOX/DMSO and DOX/DMOG treatment groups. Z’ and fold change (FC) for the Tomato/EGFP ratio for each individual plate was then calculated as per ^34^:

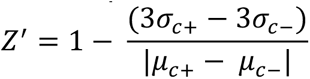

Z’ for every plate across each system was confirmed to be >0.5. Overall robustness of each system is the average of every individual Z’ and FC for each system. For temporal high content imaging, HIF, PGR and synHIF lines were seeded in plates and treated with requisite ligands immediately prior to HCI. Four treatment replicates per plate were used to assess the polyclonal population. 4 treatments per plate were used to assess the synFIH monoclone (DOX, DMSO, DOX/DMSO, DOX/DMOG), with 100ng/μL Doxycycline utilised, and 8 treatments per plate (vehicle, DMOG or R5020) were used to assess the PGR and HIF monoclonal lines. Plates were humidified and maintained at 37^0^C, 5% CO_2_ throughout the imaging experiment. Plates were then imaged every 2 hours for 40-48 hours. At every timepoint, a minimum 2000 nuclei were resampled from each well population.

### T47D mcdFLASH-PGR R5020 Dose response curve EC50 calculation

T47D mcdFLASH-PGR cells were treated with increasing doses of 0.01-100nM R5020 and quantified by HCI after 48hrs. Tomato/GFP values were min/max normalised 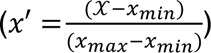 within each experiment (n = 3) and the EC50 constant and curve fitted using the drc R package from ^72^.

### Bimodal small molecule screen to identify activators or inhibitors of the hypoxic response pathway

Library of natural and synthetic compounds was supplied by Prof. Ronald Quinn and Compounds Australia, available by request. 5mM of each of the 1595 compounds were spotted in 1μL DMSO into Costar Cat#3603 plates and stored at −80°C prior to screening. Plates were warmed to 37°C prior to cell addition. Monoclonal HIF HEK293T reporter cells were seeded at 0.5×10^4^ cells per well across 20 Costar Cat#3603 plates pre-spiked with 5mM of compound in 1uL of DMSO in 100uL. On each plate, 4 wells were treated with matched DMSO amounts to compound wells as were four 1mM DMOG controls. Plates were then imaged using HCI (described above) at 36 hrs or 24 hours for reporter activation. Wells were then treated with 100uL of 2mM DMOG (for 1mM DMOG final, 200uL media final). 4 vehicle and 8 DMOG-treated controls (excluding the initial controls from the activator screen) were used for the inhibitor screen. Cells were imaged again 36 hours (Screen 1) or 24 hours (Screen 2) after treatment with 1mM DMOG in the compound wells. Data was Z scored and control wells were used to establish gating for abnormal expression of Tomato and EGFP fluorophores. For the activator screen, compounds within +/− 2SD EGFP MFI of vehicle wells were counted as having unchanged transcriptional effects. Compounds with Tomato/EGFP ratio greater than +2SD of vehicle controls was counted as a putative hit. For the inhibitor screen, compounds within +/− 2SD EGFP MFI of DMOG controls were counted as having unchanged GFP expression and Compounds with Tomato/EGFP ratio lower than −2SD from the DMOG control were considered putative inhibitors. To correct for false positives within each screen, Z scored compounds were converted to their respective P score and adjusted with a ^73^ correction. Pearson correlations were then used to compare compound expression between screens with the base R package (4.4.0). Putative activators and inhibitors identified in the screens were re-spotted at 1mM, 2.5mM and 5mM in 1μL of DMSO in Costar Cat#3603 96 well trays. Activators were rescreened by HCI after 24 hours against 1×10^4^ cells HIF reporter monoclones in biological duplicate against with vehicle and 1mM DMOG controls in 100μL. Inhibitors were rescreened by HCI after 24 hours in duplicate against 1×10^4^ cells HIF reporter monoclones with 1mM DMOG to compound wells. Final compound concentrations were 10μM, 25μM and 50μM respectively and Tomato MFI and Tomato/EGFP ratio for each compound was assessed.

### Reverse Transcription and Real Time PCR

Cells were seeded in 60mm dishes at 8×10^4^ cells per vessel overnight before treatment for 48 hours with 1mM DMOG or 0.1% DMSO. Cells were lysed in Trizol (Invitrogen), and RNA was purified with Qiagen RNAEasy Kit, DNaseI treated and reverse transcribed using M-MLV reverse transcriptase (Promega). cDNA was then diluted for real time PCR. Real-time PCR used primers specific for *HIF-1α*, and human RNA Polymerase 2 (*POLR2A*) (**Supplementary Table 4**). All reactions were done on a StepOne Plus Real-time PCR machine utilising SYBER Green, and data analysed by ‘QGene’ software. Results are normalised to *POLR2A* expression. RT-qPCR was performed in triplicate and single amplicons were confirmed via melt curves.

### Flow cytometry analysis and sorting (FACS)

Prior to flow cytometry, cells were trypsinised, washed in complete media and resuspended in resuspended in flow cytometry sort buffer (Ca^2+^/Mg^2+^-free PBS, 2%FBS, 25mM HEPES pH 7.0) for cell sorting) prior to cell sorting or flow cytometry analysis buffer (Ca^2+^/Mg^2+^ free PBS, 2%FBS, 1mM EDTA, 25mM HEPES pH 7.0) for analysis followed by filtration through a 40μM nylon cell strainer (Corning Cat#352340. Cell populations were kept on ice prior to sorting. Flow cytometry was performed either using the BD Biosciences BD LSRFortessa or the BD Biosciences FACS ARIA2 sorter within a biosafety cabinet and aseptic conditions, using an 85µM nozzle. Cell populations were gated by FSC-W/FSC-H, then SSC-W/SSC-H, followed by SSC-A/FSC-A to gate cells. EGFP fluorescence was measured by a 530/30nm detector, and the Tomato fluorescence was determined with the 582/15nm detector. A minimum of 10,000 cells were sorted for all FACS-based analysis. Data is presented as log_10_ intensity for both fluorophores. Tomato induction was gated from the top 1% of the negative control population. Cell counts for histograms are normalised to mode unless stated otherwise. FACS analysis was done on FlowJo^TM^ v10.9.1 software (BD Life Sciences)^74^.

### Time Lapse Spinning Disc Confocal Microscopy

HEK293T mcdFLASH-HIF and T47D mcdFLASH-PGR cells were seeded at 1×10^5^ or 5×10^5^ cells per dish respectively, onto 50µg/mL poly-D-lysine µ-Dish 35 mm, high Glass Bottom dishes (Ibidi, #81158) in FluoroBrite DMEM (Gibco, A1896701)/10% FBS/ 1% Pens/1% Glutamax/10mM HEPES pH7.9 and incubated overnight at 37°C with 5% CO2 prior imaging. Cells were treatment with either 0.5mM DMOG (mcdFLASH-HIF) or 100nM R05020 (mcdFLASH-PGR) immediately prior to imaging with a CV100 cell voyager spinning disk confocal Tomato (561 nm, 50% laser, 400ms exposure and 20% gain) and EGFP (488 nm, 50% laser, 400ms exposure and 20% gain) fluorescence for 48 hours post treatment with 15min imaging intervals. Images were captured at 40x with an objective lens with a ∼30μm Z stack across multiple fields of view. Maximum projected intensity images were exported to Image J for analysis and movie creation.

### Cell Lysis and Immunoblotting

Cells were washed in ice-cold PBS and lysates were generated by resuspending cells in either cell lysis buffer (20mM HEPES pH 8.0, 420mM NaCl_2_, 0.5% NP-40, 25% Glycerol, 0.2mM EDTA, 1.5mM MgCl_2_, 1mM DTT, 1x Protease Inhibitors (Sigma)) (**Supp Figure 4**) or urea lysis buffer (6.7M Urea, 10mM Tris-Cl pH 6.8, 10% glycerol, 1% SDS, 1mM DTT) (**Figure 6, Supp Figure 8, 9**). Quantification of protein levels was done by Bradford Assay (Bio-Rad). Lysates were separated on a 7.5% SDS-PAGE gel and transferred to nitrocellulose via TurboBlot (Bio-Rad). Primary Antibodies used were anti-HIF1α (BD Biosciences #), anti-HA (HA.11, Biolegend #16B12), anti-Tubulin (Serotec #MCA78G), anti-GAPDH (Sigma #G8796), anti-ARNT (Proteintech #14105-1-AP). Primary antibodies were detected using horseradish peroxidase conjugated secondary antibodies (Pierce Bioscience #). Blots were visualised via chemiluminescence and developed with Clarity Western ECL Blotting substrates (Bio-Rad).

### *In vitro* iron chelation activity assay

Chelation of iron for RQ200674 was measured by a protocol adapted from ^75^ for use in 96 well plate format. 0.1mM FeSO_4_ (50μL) and 50μL of RQ200674, Dipyridyl (positive control) or DMOG solutions were incubated for 1hr at room temperature prior to addition of 100μL of 0.25mM Ferrozine (Sigma) and incubated for a further 10 minutes. Absorbance was measured at 562nM. Chelation activity was quantified as:

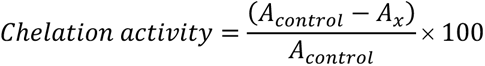

Where A_control_ is absorbance of control reactions without RQ200674, DP or DMOG and A_x_ is absorbance of solutions with compound.

## Statistical Analysis

All data in graphs were presented as a mean ± sem unless otherwise specified. Significance was calculated by a Two-Way ANOVA with Tukey multiple comparison or unpaired t-test with Welches correction where appropriate using Graphpad PRISM (version 9.0.0). All statistical analysis is from three independent biological replicates

## Figure Creation

Schematics and diagrams were created with BioRender (BioRender.com) and graphs were made either with ggplot package in R^76^ and GraphPad PRISM (version 9.0.0).

## Data Availability

Source data are provided with this paper. Additional data, including full construct sequences, are available from corresponding authors upon request. Constructs not available on Addgene can be requested from corresponding authors.

## Acknowledgements

We thank Nicholas Smith, Alexander Pace, and members of our laboratories for critical feedback and helpful discussions. We also wish to acknowledge Adelaide Microscopy and the AHMS and SAHMRI Flow Cytometry facilities for technical assistance. We acknowledge Compounds Australia (www.compoundsaustralia.com) for their provision of specialized compound management and logistics research services to the project. This work was supported by Australian Government Research Training Scholarships (T.P.A, A.E.R), The Emeritus Professor George Rodgers AO Supplementary Scholarship (T.P.A, A.E.R). The Playford Memorial Trust Thyne Reid Foundation Scholarship (A.E.R). The George Fraser Supplementary Scholarship (A.E.R), The University of Adelaide Biochemistry Trust Fund (D.J.P. and M.L.W) and the Bill and Melinda Gates Foundation Contraceptive Discovery Program [OPP1771844] (D.C.B, D.L.R).

## Author contributions

Study was initially conceived by D.C.B and M.L.W. T.P.A, D.C.B., A.E.R designed and performed experiments. T.P.A, D.C.B., M.L.W, M.L. and R.J.Q. performed and analysed the bimodal screening campaign. M.R. and A.E.R. derived FIH KO cell line. T.P.A, D.C.B and M.L.W wrote the manuscript with input from all authors. Work was supervised by D.J.P, D.L.R. & M.L.W.

## Source Data

Source data for figures is available with this manuscript.

## Correspondence and requests for materials

Should be addressed to David C. Bersten.

---

**Supplementary Table 1:** Synthetic toolkit for generation of reporter cell lines.

| **Deposit Name:** | **Availability** | **Purpose** |
| --- | --- | --- |
| **Dual fluorescent reporter constructs:** |  |  |
| pLV-REPORT(EF1a) | Addgene #172326 | Reporter with mnucTomato and EF1a downstream promoter |
| pLV-REPORT(EF1a)-TTN | Addgene #172327 | Reporter with mnucTomato-HSVtk-2A-NeoR and EF1a downstream promoter |
| pLV-REPORT(PGK) | Addgene #172328 | Reporter with mnucTomato-HSVtk-2A-NeoR and PGK downstream promoter |
| pLV-REPORT(PGK/CMV) | Addgene #172330 | Reporter with mnucTomato-HSVtk-2A-NeoR and PGK/CMV downstream promoter |
| 12xHRE-pLV-Report-EF1a | Addgene: #172333 | Reporter with HRE enhancer |
| 12xHRE-pLV-REPORT(PGK) | Addgene #172334 | Reporter with HRE enhancer |
| 12xHRE-pLV-REPORT(PGK/CMV) | Addgene #172335 | Reporter with HRE enhancer |
| PREcat-pLV-REPORT(PGK/CMV) | By Request | Reporter with a PR-responsive concatemer, with enhancers from 5 target genes, containing 6 PR response elements. |
| 5xGRE-pLV-REPORT(PGK/CMV) | Addgene #172336 | Reporter with GRE enhancer |
| 12xHRE-pLV-REPORT(EF1a) | By Request | Reporter with HRE |
| 12xHRE- pLV-REPORT(EF1a)-tdnucTomato | By Request | Reporter with tdnucTomato and EF1a downstream promoter |
| **Protein expression constructs:** |  |  |
| pLV-TET2Puro | By Request | Doxycycline-inducible expression vector |
| pLV-TET2BlastR | By Request | Doxycycline-inducible expression vector |
| pLV-TET2nucTomato | By Request | Doxycycline-inducible expression vector |
| pLV-TET2Puro-gal4DBD-miniVPR-HA | Addgene #207171 | Doxycycline-inducible expression vector for GAL4DBD-miniVPR |
| pLV-TET2Puro-gal4DBD-HIFCAD | Addgene #207173 | Doxycycline-inducible expression vector for GAL4DBD-HIFCAD (727-826) with Myc tag |
| pEF-IRES-puro6 gal4DBD-HIFCAD myc tag | Addgene #207171 | Constitutively expresses GAL4DBD-HIFCAD (727-826) with Myc tag |
| pEF-IRES-puro6 gal4DBD-HIFCAD pGalO linker | Addgene #207172 | Constitutively expresses GAL4DBD-HIFCAD (727-826) with Myc tag |
| pENTR1a-CRISPRoffv2.1 | Addgene #207174 | Lentiviral expression vector for CRISPRoffv2.1 with BFP tag |
| pLV-Egl-NeoR | Addgene #207175 | Gateway-compatible lentiviral expression plasmid with Neomycin resistance |
| pLV-Egl-BlasR | Addgene #207176 | Gateway-compatible lentiviral expression plasmid with Blastidicin resistance |
| pLV-Egl-HygroR | Addgene #207177 | Gateway-compatible lentiviral expression plasmid with Hygromycin resistance |
| pLV-Egl-ZeoR | Addgene #207178 | Gateway-compatible lentiviral expression plasmid with Zeocin resistance |

**Supplementary Table 2:**
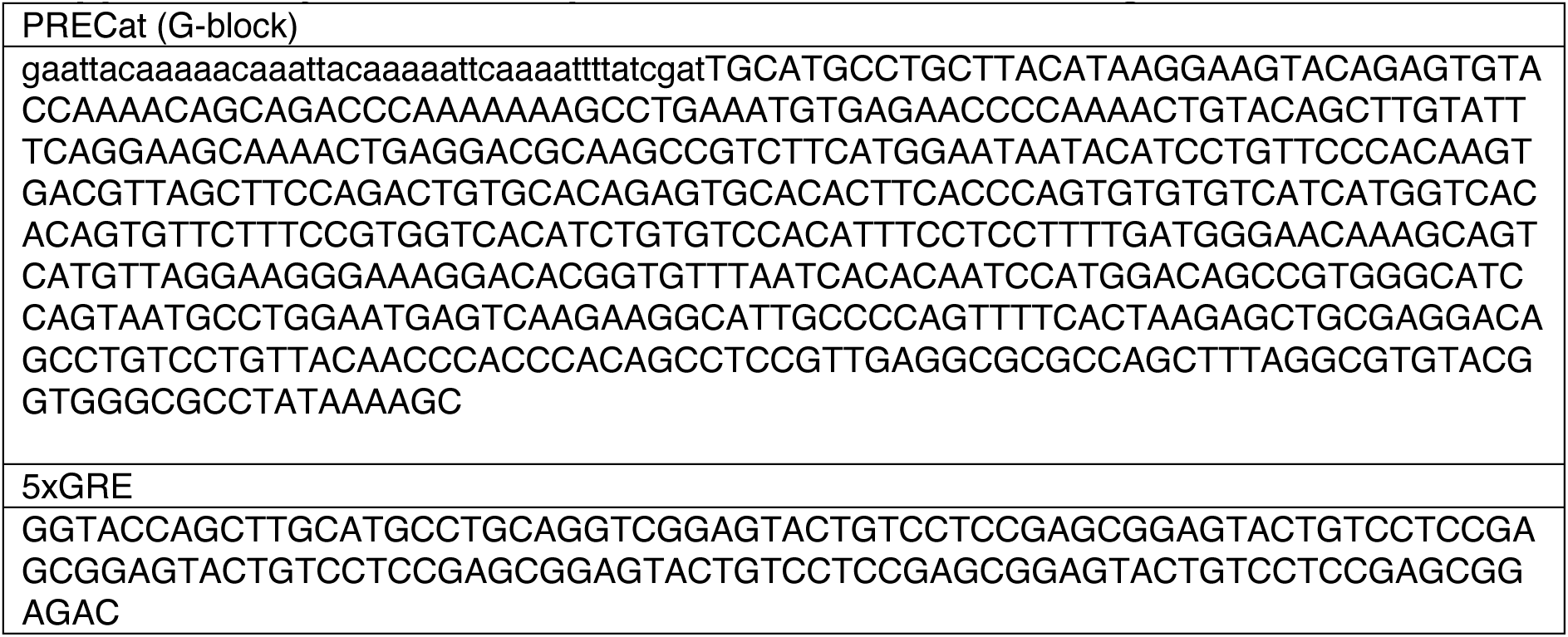
Sequences for enhancer cloning.

**Supplementary Table 3:**
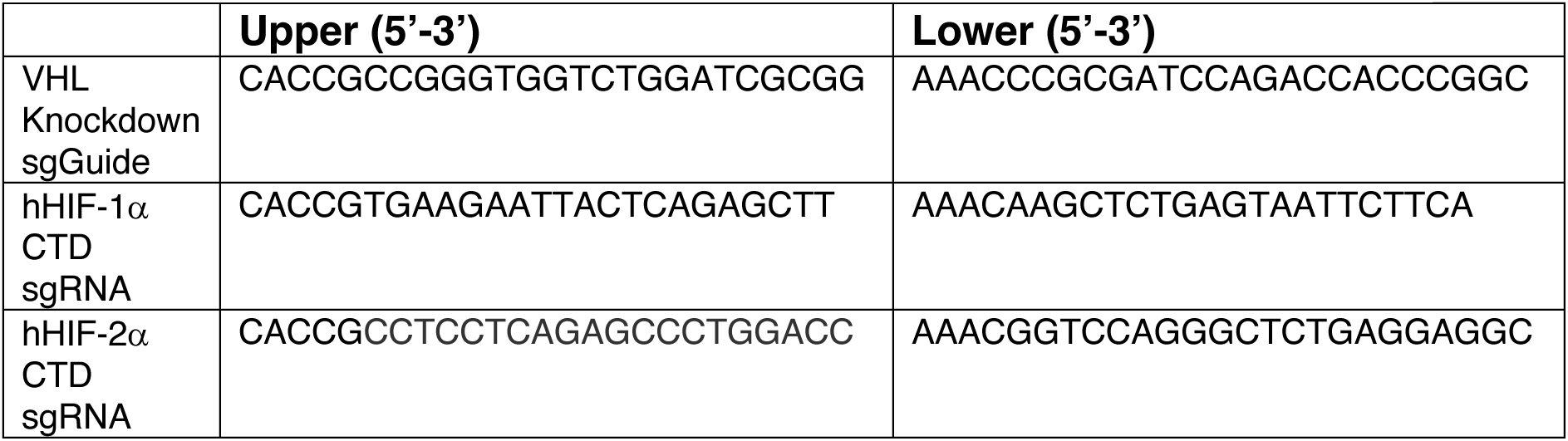
Index of all sgGuide oligos used.

**Supplementary Table 4:**
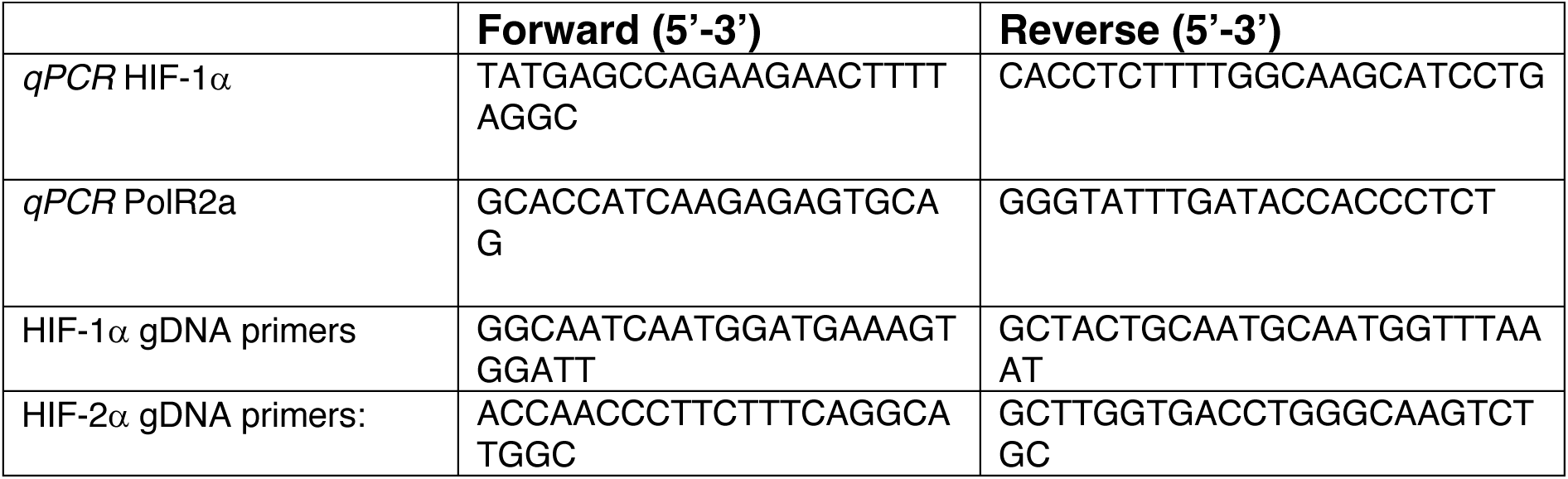
Primer sets for qPCR and PCR confirmation.

